# Effect of LSD and music on the time-varying brain dynamics

**DOI:** 10.1101/2022.04.04.486673

**Authors:** Iga Adamska, Karolina Finc

**Affiliations:** Centre for Modern Interdisciplinary Technologies, Nicolaus Copernicus University, Toruń, Poland; Faculty of Philosophy and Social Sciences, Nicolaus Copernicus University, Toruń, Poland

**Keywords:** brain states, clustering, LSD, music, psychedelics, resting-state

## Abstract

**Rationale:** Psychedelics are getting closer to being widely used in clinical treatment. Music is known as a key element of psychedelic-assisted therapy due to its psychological effects, specifically on the emotion, meaning-making and sensory processing. However, there is still a lack of understanding in how psychedelics influence brain activity in experimental settings involving music listening.

**Objectives:** The main goal of our research is to investigate the effect of music, as a part of “setting”, on the brain states dynamics after lysergic acid diethylamide (LSD) intake.

**Methods:** We used an open dataset, where a group of 15 participants underwent two functional MRI scanning sessions under LSD and placebo influence. Every scanning session contained three runs: two resting-state runs separated by one run with music listening. We applied K-Means clustering to identify the repetitive patterns of brain activity, so-called *brain states*. For further analysis, we calculated states’ dwell time, fractional occupancy and transition probability.

**Results:** The interaction effect of music and psychedelics led to change in the time-varying brain activity of the task-positive state. LSD, regardless of the music, affected the dynamics of the state of combined activity of DMN, SOM and VIS networks. Crucially, we observed that the music itself could potentially have a long-term influence on the resting-state, in particular on states involving task-positive networks.

**Conclusions:** This study indicates that music, as a crucial element of “setting”, can potentially have an influence on the subject’s resting-state during psychedelic experience. Further studies should replicate these results on a larger sample size.

## Introduction

Multiple clinical trials are investigating the therapeutic potential of psychedelics in treatment of various psychiatric diseases, such as depression, end-of-life anxiety, post-traumatic stress disorder, and addiction. As we are getting closer to widely approved psychedelics-assisted therapies, there is a necessity to understand the dynamical brain processes that underlie psychedelic experience in various treatment settings. Factors known as “set” and “setting” are one of the key factors modulating the effects of psychedelics and contributing to the positive outcomes of psychedelic experience (Carhart-Harris et al., 2018). “Set” refers to the individual: the person’s personality, expectations, and motivations for the session. In turn, “setting” is defined as the environment of the session. This does not only mean the exact site where the session takes place, but also the people who accompany the person together with any additional elements such as music, plants, photographs, pictures, etc. (Eisner, 1997). Here, we investigate how music, as a part of the “setting”, affects the brain states’ dynamics during the psychedelic experience.

Psychedelics, such as psilocybin, lysergic acid diethylamide (LSD) or MDMA, cause a profound effect on perception (Kometer et al., 2015; Kraehenmann et al., 2017), consciousness (Lebedev et al., 2015; Liechti et al., 2017; Smigielski et al., 2019), mood (Forstmann et al., 2020; Schmid & Liechti, 2018), personality (Lebedev et al., 2016), and behavior (Griffiths et al., 2018; Kometer et al., 2012; Schmid & Liechti, 2018). There is growing evidence that psychedelic substances can increase openness (MacLean et al., 2011), enhance creative thinking, along with empathy and subjective well-being (Mason et al., 2019; Pokorny et al., 2017). In addition, altered states of consciousness, which contribute to the “mystical-type experiences” phenomena, seem to be one of the most important factors that can cause positive outcomes of the psychedelic experience (Barrett & Griffiths, 2017; Griffiths et al., 2006, 2011, 2018; Liechti et al., 2017; MacLean et al., 2011). At present, psychedelic-assisted therapy finds use in treating depression (Carhart-Harris, Bolstridge, et al., 2016; Davis et al., 2021), cancer-related psychological distress (Gasser et al., 2015; Griffiths et al., 2016; Grob et al., 2011; Ross et al., 2016), addiction (Bogenschutz et al., 2015; Bogenschutz & Johnson, 2016; Johnson et al., 2014) and post-traumatic stress disorder (Mithoefer et al., 2019; Ot’alora G et al., 2018; Varker et al., 2021). What can contribute to better targeting and personalization of psychedelic-assisted therapies is the increased understanding of the exact mechanisms of their action on the human body, specifically, the brain.

Acting mostly on serotonergic 5-HT_2A_ receptors and, to lesser extent, on dopaminergic and glutamatergic neuromodulatory systems, psychedelics influence the whole-brain activity and network organization. Research on the psychedelics influence on resting-state found that they can modulate the exploration of the brain’s repertoire of functional network states (Lord et al., 2019; Tagliazucchi et al., 2014), alter their segregation and integration (Luppi et al., 2021; Preller et al., 2020), increase whole-brain between-network functional connectivity (Mason et al., 2020), decrease modularity of large-scale brain networks (Lebedev et al., 2015) and support formation of new local-range connections (Petri et al., 2014). In terms of specific brain subsystems, studies report an decrease in activity in medial prefrontal cortex (mPFC), a key region in default mode network (DMN) (Carhart-Harris, Erritzoe, et al., 2012; Palhano-Fontes et al., 2015), along with decreased functional connectivity within DMN (Carhart-Harris, Erritzoe, et al., 2012; Carhart-Harris, Muthukumaraswamy, et al., 2016). Even though studies have shown evidence for recurrent states in resting-state (Cabral et al., 2017; Cornblath et al., 2020; Vidaurre et al., 2017), it was Singleton et al. (2022) who also showed that the brain manifests recurrent states of brain network activity under LSD influence. Since the majority of neuroimaging research on psychedelics focused on comparing brain activity or connectivity at rest, we still need research on how psychedelic experience affects the brain dynamics during different experimental settings.

Music has been an inherent part of spiritual ceremonies for centuries (Guerra-Doce, 2015) and appears to be the most frequently used stimuli during psychedelic sessions (O’Callaghan et al., 2020). As a part of the “setting” (Eisner, 1997), its role is to enhance and influence the sensory and mental effects of psychedelics as well as serve as a “guide” supporting the whole experience (Bonny & Pahnke, 1972). Music can deeply affect human cognition, therefore, its potential to contribute to the positive outcomes of the psychedelic-assisted therapy should not be underestimated (Barrett et al., 2017; Barrett, Preller, & Kaelen, 2018). Indeed, recent studies confirm that music supports the beneficial effect of psychedelics on mental imagery (Kaelen, Roseman, Kahan, et al., 2016), emotion processing (Carbonaro et al., 2018; Kaelen et al., 2015; Kaelen, Roseman, Lorenz, et al., 2016), meaning making (Preller et al., 2017) and openness (Kaelen et al., 2018). However, most studies focus on the psychological aspect of using music in psychedelic therapy, rather than investigate its impact on the brain network organization and fluctuations of brain activity in time. As a result, brain activity under music and psychedelic influence remains poorly understood. An important question remains: how both of these factors facilitate changes in brain architecture? Can we also observe changes in time-varying brain activity during psychedelic experience combined with music listening?

Our goal was to examine the effect of LSD and music on time-varying brain dynamics. The dynamics of time-varying brain activity can be captured by determining *brain states*, that is, repetitive patterns of cortical activity (Cornblath et al., 2020; Meer et al., 2020). In detail, brain states represent the whole-brain activity at a given time point, which occur repeatedly during certain condition and are reproducible across subjects. The major advantage of this method, in comparison to standard brain activation analysis methods based on General Linear Model (Friston et al., 1994), is the ability to focus more on the temporal rather than spatial resolution of the data, which makes it suitable to track changes in brain activity during psychedelic experiences, however there are still limitations which result from the speed of the hemodynamic fluctuations and low temporal resolution of the fMRI. Here, we applied the K-Means clustering algorithm (Allen et al., 2014; Cornblath et al., 2020; Goutte et al., 1999; Le Cam & Neyman, 1967; Lloyd, 1982) to identify brain states and to compare their dynamics during resting-state and music listening. We used an open functional MRI dataset from the intervention study, where a group of 15 participants underwent two scanning sessions under LSD influence and under placebo. Every scanning session contained three runs: two resting-state runs separated by one run with music listening.

First, we compared resting-state without (1st run) and with music (2nd run) stimuli during psychedelic experience and on placebo, to investigate how music affects brain states dynamics. Based on previous studies on music processing (Blood & Zatorre, 2001; Chan & Han, 2022; Koelsch, 2011; Vuilleumier & Trost, 2015) we hypothesized that brain states associated with emotions (limbic network), autobiographical memory (default mode network), and music processing (sensory networks) will be more frequent during music listening in comparison to resting-state. Second, we investigated the long-term effect of music under LSD by comparing brain states dynamics during resting-state *before* (1st run) and resting-state *after* music listening (3rd run). Since music is an essential element supporting positive outcomes of psychedelic treatments (Barrett et al., 2017; Kaelen et al., 2018), we hypothesized that music listening on psychedelics will have a lasting impact on the time-varying brain activity. Specifically, we expect to observe different patterns of brain states dynamics for the two resting-state runs. Psychedelics are known to enhance sensory (Aday et al., 2021; Carhart-Harris, Erritzoe, et al., 2012; Carhart-Harris, Muthukumaraswamy, et al., 2016; Császár-Nagy et al., 2019), emotional (Grimm et al., 2018; Kometer et al., 2012; Kraehenmann et al., 2015) and self-related processing (Smigielski et al., 2020) therefore, in both cases we hypothesize that besides music, LSD may be also an important factor in affecting the brain states dynamics. LSD was reported to have an effect on the brain entropy by increasing it in the sensory and higher networks (Lebedev et al., 2016) and here its effects could be manifested by making the states’ transitions more irregular and frequent. Additionally, the appearance of states related to autobiographical memory and self-focused stimuli could be more pronounced, whether by a longer duration time or by higher appearance rate in comparison to other states.

## Methods

### Dataset

Data used for analysis was obtained from the OpenNeuro database (OpenNeuro Dataset ds003059) (al., 2020; Carhart-Harris, Muthukumaraswamy, et al., 2016). Dataset included preprocessed fMRI scans obtained from 15 participants in two sessions, each consisting of three runs. Detailed recruitment of subjects is described in Carhart-Harris et al. (2016).

### Experimental procedures

Full description of the experimental procedures can be found in Carhart-Harris et al. (2016). In summary, twenty healthy participants were recruited for two sessions of fMRI scanning. Sessions were separated by at least 14 days. In one of these sessions, subjects received LSD (75 μg in 10 mL saline) and in the other placebo (10 mL saline). Following the post-infusion acclimatization period, fMRI scanning started 115 minutes after the infusion. Each session consisted of three runs, lasting approximately 7.5 minutes. During each scan participants were encouraged to keep their eyes closed and relax. First and third runs were resting-state, while the second involved listening to music (Fig. 1). Two tracks from the album Yearning, made by ambient artist Robert Rich and classical Indian musician Lisa Moskow were chosen for listening during this run. Each participant listened to both tracks, in a balanced order across conditions (Kaelen, Roseman, Kahan, et al., 2016). For data analysis we included fMRI images from all runs, as our main goal was to investigate both the effect of LSD and music on brain states’ dynamics.

**Fig. 1.**
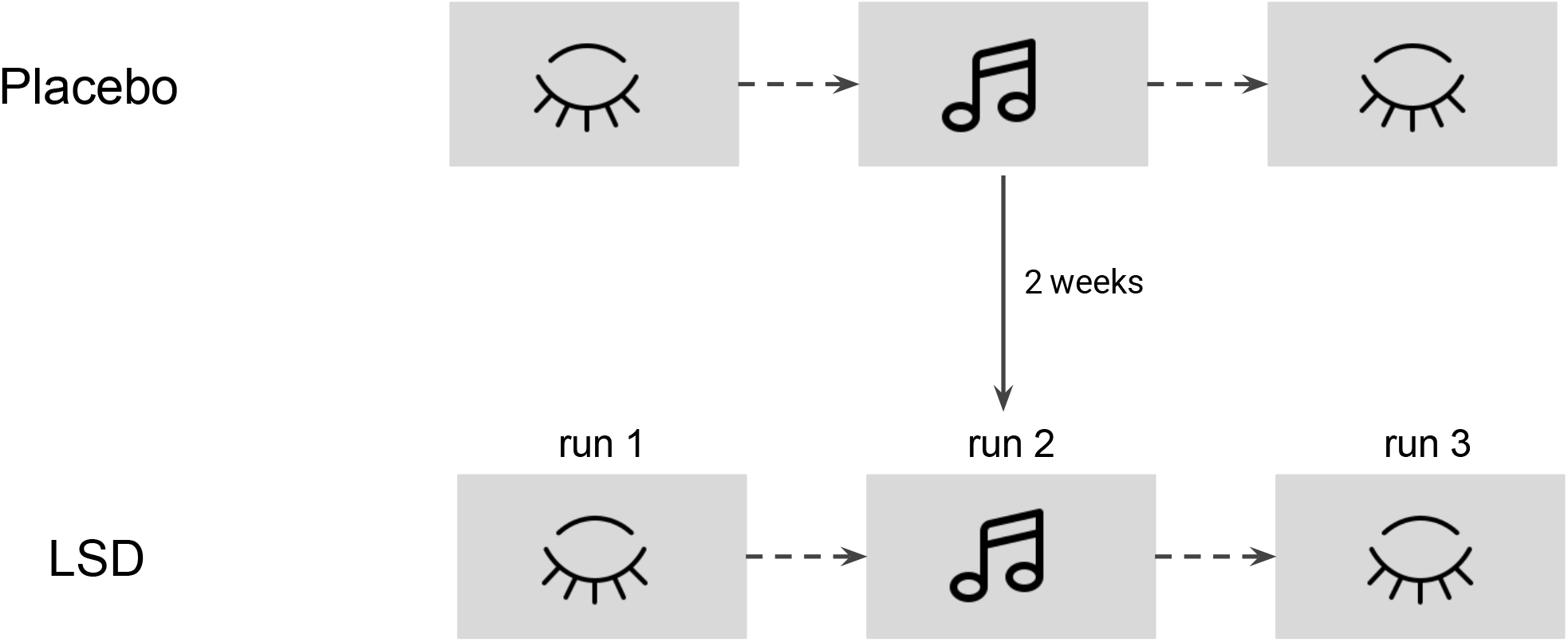
Study design. Participants underwent two single-blind fMRI sessions in either LSD or placebo condition. The order of conditions was balanced across subjects. Each session included three scans, 1st and 3rd with no stimulation (resting-state) and the 2nd, which involved listening to music. Each run lasted approximately 7.5 minutes. Musical Notes, Closed eye icon by Icons8.

### Data acquisition and preprocessing

The detailed overview of data acquisition and preprocessing is described in Carhart-Harris et al. (2016). In brief, BOLD fMRI data acquisition was performed using a gradient echo planar imaging sequence, with TR = 2 s, FOV = 220 mm and 64 × 64 acquisition matrix. Preprocessing of the BOLD fMRI data consisted of: 1) removal of the first 3 volumes; 2) despiking (AFNI); 3) slice-time correction (AFNI); 4) motion correction (AFNI); 5) brain extraction (BET, FSL); 6) registration to anatomical scans (FSL, Freesurfer, manual); 7) non-linear registration to 2mm MNI brain (Symmetric Normalization (SyN), ANTS); 8) scrubbing; 9) spatial smoothing (FWHM) of 6mm (AFNI); 10) band-pass filtering between 0.01 to 0.08 Hz (AFNI); 11) de-trending (AFNI); 12) regression of the 9 nuisance regressors: 3 translations, 3 rotations, 3 anatomically-related (not smoothed) (Symmetric Normalization (SyN). After the preprocessing procedures 15 and 12 participants were included for analysis, from resting-state and music runs respectively. Only 12 participants qualified for the analysis since 3 had technical problems with music.

### Brain parcellation

For signal extraction, we used Schaefer 2018 atlas parcellation, with 400 ROIs and 2 mm resolution (Schaefer et al., 2018). The parcellation was downloaded from a publicly available repository (https://github.com/ThomasYeoLab/CBIG/tree/master/stable_projects/brain_parcellation/Schaefer2018_LocalGlobal, commit: 703454d).

### Clustering of BOLD scans using KMeans algorithm

In order to represent brain activity and its fluctuations as so-called brain states (Cornblath et al., 2020) we applied the K-Means clustering algorithm (Allen et al., 2014; Cornblath et al., 2020; Goutte et al., 1999; Hartigan & Wong, 1979; Le Cam & Neyman, 1967; Lloyd, 1982). K-Means is an unsupervised machine learning method that allows to group data points with similar spatial properties with no prior knowledge about the structure of the data. Here, brain states are defined as repeated patterns of brain activity, represented as clusters in the regional activation space. Prior to clustering timeseries into discrete brain states, we concatenated all data points into one *N* × *P* array, where *N*-number of timepoints, *P*-number of features (ROIs). The length of *N* was equal to 18228 (15 subjects × 868 resting-state scans + 12 subjects × 434 music scans). For the final concatenated timeseries used in clustering, we included data from both LSD and placebo sessions and from both conditions (resting-state and music listening). Such a procedure ensures the correspondence of brain states’ labels across all subjects, sessions and runs To determine the most appropriate number of brain states, we applied the K-Means algorithm with Euclidean distance as a measure of similarity. Following Cornblath et al. (2020), we identified the maximum number of *k*. For both conditions (LSD and placebo) the length of timeseries was equal to 217, therefore to analyze the transitions between states based on the *k* × *k* matrix, *k*^2^ must be < 217. Based on that, we run the K-Means clustering with *k* ranging from 2 to 14. We repeated this procedure 100 times, with random seed = 42 to ensure reproducibility. For the main criterion for choosing the optimal *k*, we selected the gain in variance explained for a unit increase in *k*. Variance explained is defined as the ratio of between-cluster variance (sum of squares) to total variance in the data (within-cluster plus between-cluster sum of squares) (Cornblath et al., 2020). As shown in Supplementary Fig. 2, for *k* >5 gain in variance explained is equal to less than 1%, therefore we have chosen *k* = 4 as the most optimal number of brain states (Cornblath et al., 2020). These results are consistent with Singleton et al. (2022), who also used the K-Means algorithm to cluster resting-state scans of this dataset and also found *k* = 4 as the most appropriate number of *k*. Additionally, we checked for the number of states for each subject and each *k*, as there is a possibility that some subjects will not manifest all states (some will be absent). We estimated the number of absent states by calculating the sum of absent appearances for each number of states and plotted them for each run and session. The analysis showed that for *k* >13 the correspondence of brain states across participants was not maintained (Supplementary Fig. 3). It is important to note that selecting the optimal number of states based on the gain in variance explained is one of the possible options of choosing *k*. One could hypothesize that a higher number of brain states could provide a presentation of brain dynamics at different levels, possibly representing changes on a smaller temporal scale. Therefore, we replicated all analyses for *k* = 5 and for *k* = 6. The results are provided in Supplementary Information.

### Correlating brain states with large-scale brain networks and Neurosynth topic maps

After the KMeans clustering, we correlated each brain state with 7 a priori defined brain networks from the Schaefer 2018 atlas parcellation. The parcellation was provided together with labels that indicated the membership of each parcel to one of the 7 large-scale networks. As a result, each brain state activity could be associated with positive or negative correlation with each large-scale brain network, as shown on Fig. 2. Additionally, to get more insight into the brain states’ characteristics we correlated each state with several uniformity test maps obtained from the Neurosynth database (https://neurosynth.org/). Uniformity test maps are defined as z-scores from a one-way ANOVA testing whether the proportion of studies that report activation at a given voxel differs from the rate that would be expected if activations were uniformly distributed throughout gray matter. Here, we correlated each brain states’ spatial distribution to 11 uniformity test maps which can potentially describe the psychedelic experience: social cognition, attention, music, emotional, somatosensory, happy, pain, reward, language, anxiety and autobiographical memory. This procedure allowed us to describe each state in terms of associated mental states (Fig. 3).

**Fig. 2.**
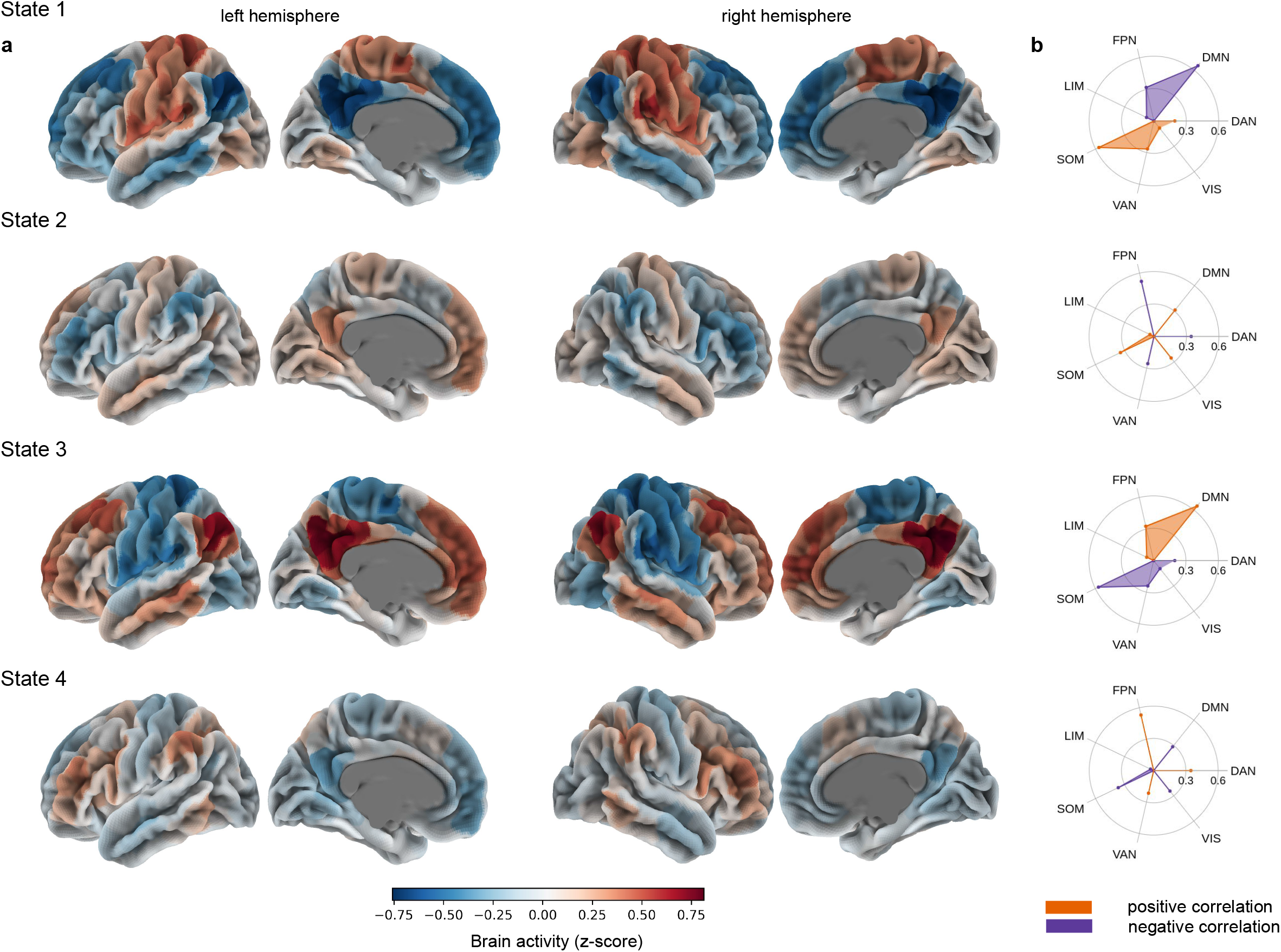
Brain states’ correlation with large-scale brain networks. **a** Each state is defined as a characteristic pattern of brain activity, which fluctuates from high to low. **b** Correlation with 7 large-scale brain networks obtained from Schaefer 2018 parcellation (FPN - frontoparietal network, DMN - default mode network, DAN - dorsal attention network, VIS - visual network, VAN - ventral attention network, SOM - somatomotor network, LIM - limbic network).

**Fig. 3.**
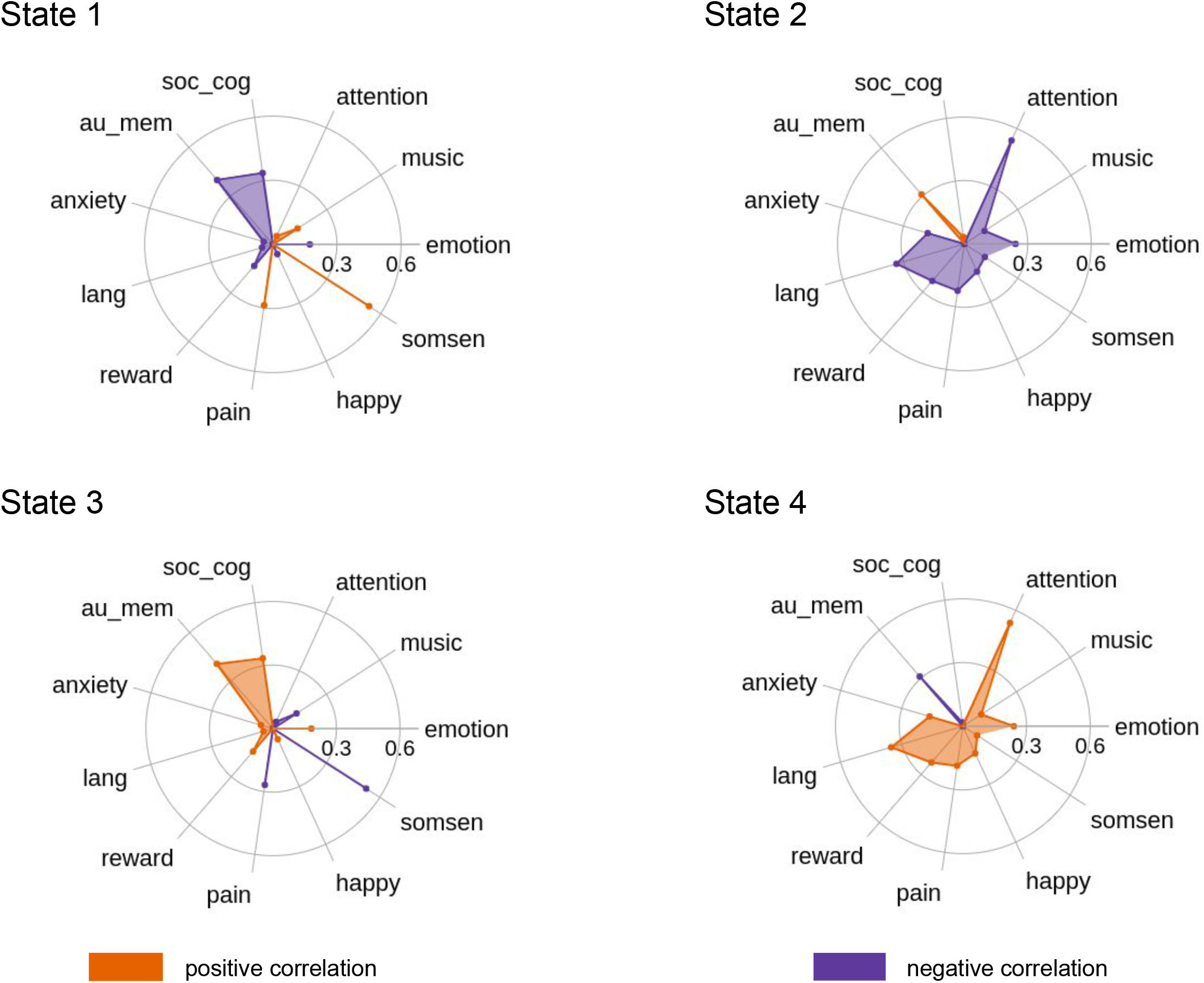
Brain states’ functional profiles. Uniformity test maps obtained from the Neurosynth database can potentially describe each brain state in the terms of potentially associated mental states, which can occur during psychedelic experience (soc_cog - social cognition, emotion - emotional, somsen - somatosensory, lang - language, au_mem - autobiographical memory).

### Analysis of the time-varying brain activity

Next, we calculated three measures that allow us to investigate and analyze dynamics of the brain states: fractional occupancy, dwell time, and transition probability. Fractional occupancy is defined as the number of scans classified as the particular state, divided by the total number of scans. We can interpret the fractional occupancy measure as a time spent in a particular brain state relative to the total time spent during a run. Dwell time is calculated as the number of all time points classified as a particular state divided by the number of their consecutive appearances (eg. 11122112221; for state 1 dwell time is 2 since there are 6 time points in total which appear in 3 consecutive appearances); it represents the mean length of a duration of each state. Transition probability is defined as the probability of transitioning from state *i* to state *j*. The measure allows us to infer what was the common transition pattern between all brain states. Each measure was calculated on a subject level.

### Statistical analysis and reproducibility

The main goal of our analyses was to investigate the effect of music on brain state activity after LSD intake. To approach this goal, we performed two major statistical comparisons. First, between resting-state prior to music listening (1st run) and music listening (2nd run), and second between both resting-state sessions (pre- and post-music experience; 1st and 3rd run) (Fig. 1). To check whether the analyzed variables (fractional occupancy, dwell time and transition probability) are normally distributed we performed a series of Kolmogorov-Smirnov tests. In each analysis, that is for comparison of the resting-state with music listening and resting-state *before* and *after* listening to music, for each of the four states we used a two-level multilevel model (MLM). The MLM analysis was performed using lmer and emmeans R package. The model was specified with either fractional occupancy or dwell time as the dependent variable and with session (two factors: placebo and LSD) and run (two factors: run 1 and run 2 or 3, depending on the analysis) as independent variables. In addition to the main effects (session, run) we also included the session × run interaction term. In all models, we specified fixed effects for session, run and interaction and random effects for individuals. Such a model allows subjects to vary in terms of their intercept and their effect of session and run (random slopes). Since the transition probability measures did not meet the criteria of normal distribution, we used the permutation non-parametric test to perform statistical analyses for both LSD and placebo sessions. In the permutation test we used the mean difference between two conditions, with 10000 permutation samples and random seed = 0. Transition probabilities of both experimental (LSD session) and control (placebo session) data were shuffled randomly, within subjects. To test the significance of the permutation test we applied t-statistics. We set up the level of alpha to 0.05 in all analyses. Due to the small group sample (15 participants in resting-state run, 12 in music listening run), thus a low power of the study, t-test results were not corrected for multiple comparisons. To ensure reproducibility, Python and R code for all analyses is publicly available (see section “Code availability”). Note that these analyses were performed on a small sample, therefore the results should be interpreted with caution and considered as preliminary.

### Code availability

All code scripts used for analyses can be found at: github.com/igaadamska/LSD-music-brainstates

## Results

### Brain states characteristics

To investigate changes in the brain signal dynamics and analyze them under the influence of music listening and psychedelic experience, we identified repeated patterns of brain activity — *brain states*. To cluster fMRI timeseries into brain states, we used the K-Means clustering algorithm, which assigns each data point to the nearest of the a priori defined centroids (Hartigan & Wong, 1979; Le Cam & Neyman, 1967; Lloyd, 1982). After calculating K-Means’ measures (within- and between-cluster sum of squares, silhouette score, variance explained, gain in variance explained; Supplementary Fig. 1, 2), we chose *k* = 4 as the most optimal number of centroids (see “Methods”).

The clustering analysis revealed four brain states, each of which can be characterized by a specific pattern of cortical activity of 7 a-priori defined networks (see “Methods”). State 1 is described by the high activity of the somatomotor (SOM) network and low activity of the default mode (DMN) and frontoparietal (FPN) networks (Fig. 2). Its activity pattern highly corresponds to the activation of the primary brain regions in response to basic visual, motor or tactile stimuli. Additionally, correlation with Neurosynth topic maps revealed that this state can manifest during music listening or pain perception (Fig. 3). State 2 is associated with the combined average activity of DMN, SOM and VIS networks and low activity of task-positive networks (Fig. 2). This pattern of activation could be associated with self-referential and simple sensory stimuli processing; VIS networks activity may reflect the processing of visual stimuli, which is important in the context of the psychedelic experience for processing hallucinations (Aday et al., 2021; Johnson et al., 2019; Kometer, Schmidt, Jancke, et al., 2013). Interestingly, based on the Neurosynth topic maps analysis, combined activity of these three networks could be crucial for autobiographical memory processing, similarly to state 2 (Fig. 3). State 3 presents the opposite activation pattern to state 1, which is characterized by a high activity of the DMN and the FPN, along with a low activity of the SOM network (Fig. 2). High DMN activity suggests that this state might be related to self-referential processing or mind-wandering, that are often altered after psychedelics intake (Carhart-Harris, Erritzoe, et al., 2012; Carhart-Harris, Leech, et al., 2012; Johnson et al., 2019; Kometer et al., 2015). In line with that, Neurosynth correlation analysis showed a strong similarity of this state activity to activity patterns occurring during social cognition and autobiographical memory (Fig. 3). Moreover, simultaneous activity of the DMN and the FPN could be also relevant to working memory performance (Cocchi et al., 2013). Finally, state 4 is described by the high activity of task-positive networks, that is, FPN, DAN and VAN (Fig. 2). Such brain activity is characteristic for facilitating cognitively demanding tasks (cognitive control), executive functions and attention, together with integration of sensory, emotional, and cognitive information (Downar et al., 2000; Vossel et al., 2014). Here, this state could be responsible for integrating different aspects of the psychedelic experience. Results revealed by Neurosynth correlation analysis showed strong correlation with activity pattern occurring during attention-demanding tasks, what is consistent with the state’s characteristics (Fig. 3).

### Effect of LSD on brain states during music experience

The multilevel linear modeling for fractional occupancy revealed that the main effects as well as interaction effects were not statistically significant for all states (Fig. 4a). In contrast, the MLM analysis for dwell time showed a significant session effect for state 2. Specifically, the dwell time for this state was significantly lower during the LSD session (*F*_1,25_ = 4.9953, *p* = 0.035) (Fig. 4b). Additionally, we found a significant session × run interaction for state 4 (*F*_1,25_ = 4.9642, *p* = 0.035) (Fig. 4b). The post-hoc analysis showed a significant difference for run 2 between the LSD and placebo sessions (paired t-test, *t* = −2.219, *p* = 0.036).

**Fig. 4.**
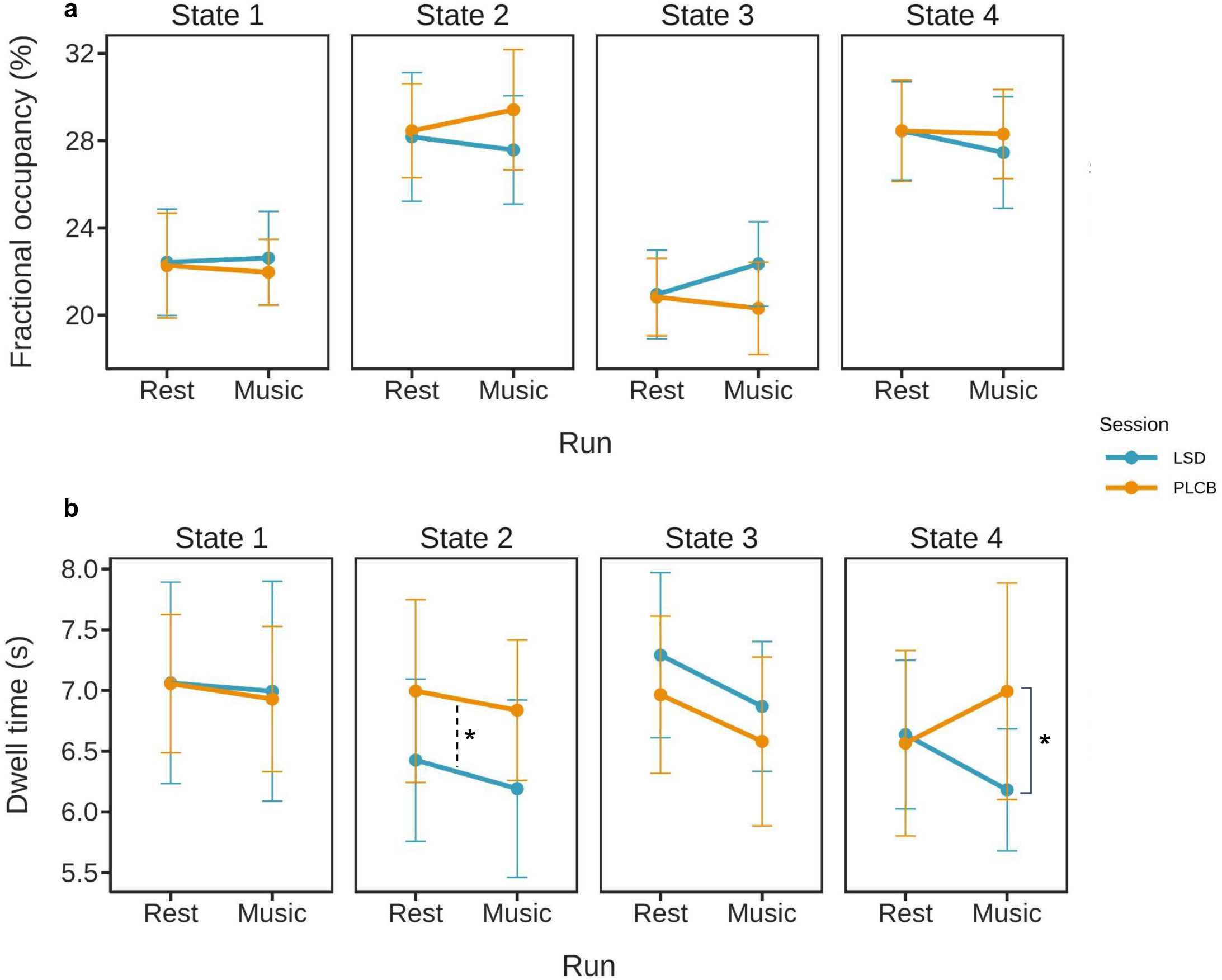
Comparison of the resting-state and music experience. **a** Fractional occupancy for 1st (resting-state) and 2nd (music listening) runs for both LSD and placebo conditions. For every state, we found no significant main and interaction effects. **b** Dwell time for 1st (resting-state) and 2nd (music listening) runs for both LSD and placebo conditions. We found a significant session effect for state 2 (*F*_1,25_ = 4.9953, *p* = 0.035) together with significant session × interaction effect for state 4 (*F*_1,25_ = 4.9642,*p* = 0.035). For state 4, music listening on LSD was associated with a lower dwell time in comparison to music listening on placebo (paired t-test, *t* = −2.219, *p* = 0.0362).

Subsequently, we were interested in how psychedelic experience combined with music can influence frequency of switches between brain states. To characterize the brain states’ dynamics we analyzed changes in transition probability using the non-parametric permutation test (see ‘Methods’). To assess the difference in states’ transitions between music listening and resting-state for each session, we performed two comparisons. In the first step, we compared transition probability between the 2nd and 1st run for the placebo session (Supplementary Fig. 5a). Secondly, we performed the same analysis for the LSD session (Supplementary Fig. 5b). For both analyses none of the results was statistically significant. Next, we wanted to track changes in transition probability that can occur during music listening between LSD and placebo (Supplementary Fig. 5c). Also here none of the results met the statistical significance criteria. Finally, we assessed whether the difference between both runs (2nd run - 1st run) could be significant (Supplementary Fig. 5d), however, the statistical analysis revealed that there is no effect. Due to small sample size and lack of correction for multiple comparisons, these results should be interpreted with caution.

### Effect of LSD and music listening on brain states during resting-state

To investigate whether brain states’ dynamic during resting-state *after* the music experience is similar to the one *before* music listening, we analyzed fractional occupancy, dwell time and transition probability data from 15 participants from both placebo and LSD sessions involving 1st run (resting-state *before* music listening) and 3rd run (resting-state *after* music listening) (Fig. 1). To investigate possible changes between resting-state periods throughout placebo and LSD conditions, we used a multilevel linear model for fractional occupancy and dwell time analysis. The analysis for all four states showed that there were no significant main and interaction effects for both of these measures (Supplementary Fig. 6a, b).

Our next step was to explore changes in brain states’ dynamics with focus on their transitions. For this reason, we used a nonparametric permutation test to analyze subjects’ transition probability (see “Statistical analysis and reproducibility”). Firstly, we compared transition probabilities of each state between 3rd run and 1st run within each session (placebo and LSD) (Fig. 5a, b). The results revealed that during LSD condition participants had higher probability of transitioning from state 3 to state 4 during the resting-state after music listening (permutation test, mean difference = 0.041, *p* = 0.024) (Fig. 5b). Additionally, we wanted to examine whether post-music listening resting-state was different on LSD in comparison to placebo. However, the mean difference was not statistically significant for any of the possible transitions. Ultimately, we performed a permutation test between differences in both runs (3rd run - 1st run) between the LSD and placebo. We found that the difference between both runs was significantly higher in the LSD session in comparison with placebo for the transition from state 3 to state 4 (permutation test, mean difference = 0.057, *p* = 0.026) (Fig. 5d). Here again, due to small sample size and lack of correction for multiple comparisons, these results should be interpreted with caution.

**Fig. 5.**
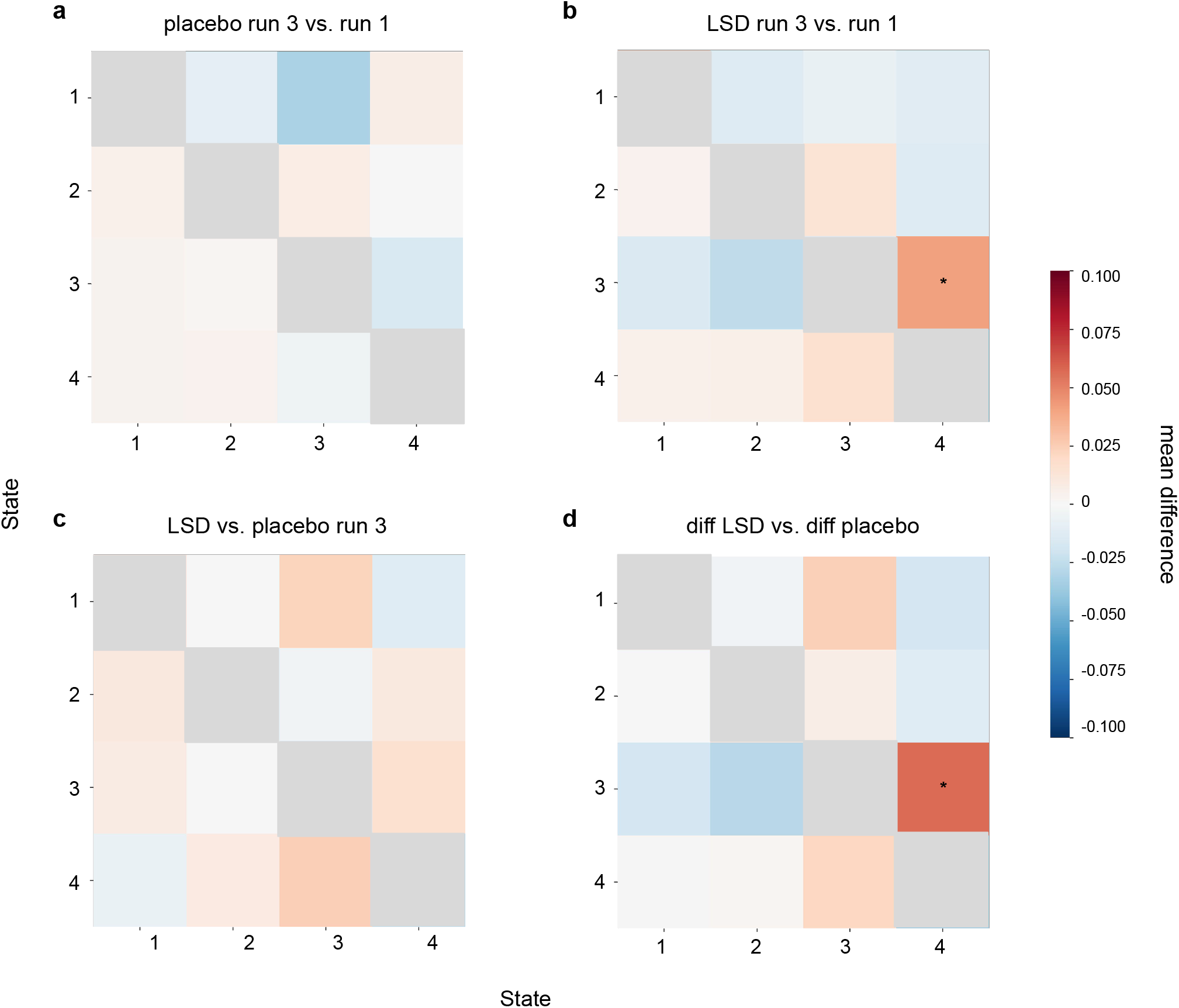
Comparison of the pre- and post-music listening resting-state. **a** Transition matrix for the placebo session, 3rd run vs. 1st run; we found no significant differences. **b** During LSD session, resting-state after music listening was associated with higher probability of transitioning from state 3 to state 4 (permutation test, mean difference = 0.041, *p* = 0.024). **c** Transition matrix for the 3rd run (resting-state after music listening), LSD vs placebo session; we found no significant differences. **d** Transition matrix for the differences (3rd run - 1st run) between both sessions (LSD and placebo). We found that for transition from state 3 to 4 the difference between both resting-states was greater after the LSD intake (permutation test, mean difference = 0.057,*p* = 0.026). For all transition matrices, the direction of transition from one state to the other is row to column.

## Discussion

In this study, we aimed to investigate the effect of LSD and music on the dynamics of time-varying brain activity. Our analysis revealed that for both sessions (LSD and placebo) and conditions (resting-state and music) the brain activity can be reduced to four repetitive patterns, so-called *brain states*. Each state is then characterized by a unique activity pattern of 7 a-priori defined brain networks and manifests different degrees of similarity to the mental states associated with psychedelic experience (see Fig. 2, 3). These four brain states can be also interpreted as two Meta-States, each of which has two sub-states which show the opposite patterns of activation (Singleton, Luppi, et al., 2022). We observed that LSD, regardless of the presence of musical stimuli, had an impact on the dynamics of the state characterized by average activity of DMN, SOM and VIS networks (state 2). Importantly, we show that the state of high activity of task-positive networks was the one most affected by the interaction between the psychedelic and music in both short-(change in mean state duration) and long-term (different transition pattern). Collectively, these findings suggest that the whole-brain states’ dynamics during psychedelic experience and placebo differs significantly and music, as a crucial element of setting, can potentially have an influence on the subject’s resting-state.

Based on previous findings on the music processing (Blood & Zatorre, 2001; Chan & Han, 2022; Koelsch, 2011) and its role during the psychedelic experience (Barrett, Preller, & Kaelen, 2018; Bonny & Pahnke, 1972), we hypothesized that music listening on the LSD will be associated with greater manifestation of brain states characterized by the high DMN, SOM, and LIM networks activity. Previous studies revealed that such network activity may be crucial for processing sensory and self-related stimuli that are often enhanced after psychedelics intake (Griffiths et al., 2006; Kometer et al., 2015; Kometer, Schmidt, Jäncke, et al., 2013; Lebedev et al., 2015; Liechti et al., 2017; Tagliazucchi et al., 2016). Here, we observed that joint effect of music and psychedelics resulted in lower dwell time, and in consequence, in lower manifestation of state 4, associated with high activity of task-positive networks in the LSD session compared to placebo. Task-positive networks, such as frontoparietal, dorsal- and ventral-attention networks are responsible for cognitive control along with integration of sensory and cognitive information (Cocchi et al., 2013; Downar et al., 2000; Vossel et al., 2014). In line with that, these results can be explained in the psychological context of the psychedelic experience: under the psychedelics influence the subject’s consciousness is more altered than during placebo and in combination with music there is a possibility that the majority of cognitive resources is used for processing the whole psychedelic session including music-induced experiences while the higher cognitive processes are suppressed. Interestingly, we found that LSD had an influence on brain states’ dynamics involving the activity of DMN, SOM and VIS networks, specifically by decreasing its mean duration. Müller et al. (2018) reported that acute LSD administration significantly decreased functional connectivity *within* visual, sensorimotor, auditory networks, and the default mode network. Therefore, we could hypothesize that these changes in the functional connectivity may also manifest when looking at recurring brain states’ activity, in particular in the decrease of dwell time of state 2. Both the diminished manifestation of this state and the fact that LSD had an effect independent of the musical stimuli make these results inconsistent with our hypothesis. Nevertheless, it supports the assumptions of LSD having the effect on time-varying brain activity. Inconsistency of our results with the previous studies may be explained by the specificity of the analysis method we used. GLM-based methods represent differences between conditions as contrasts in whole-brain activity, whereas network neuroscience tools allow to investigate changes in structural and functional brain networks. The usage of both of these approaches in the context of psychedelic experience revealed changes in the BOLD signal and whole-brain network reorganization (Lebedev et al., 2015; Lord et al., 2019; Luppi et al., 2021; Petri et al., 2014; Preller et al., 2019, 2020; Tagliazucchi et al., 2014; Varley et al., 2020), especially regarding the DMN and the visual regions (Carhart-Harris, Erritzoe, et al., 2012; Carhart-Harris, Muthukumaraswamy, et al., 2016; Palhano-Fontes et al., 2015). However, none of these approaches takes full advantage of the temporal resolution provided by the neuroimaging methods. Here, we applied the KMeans algorithm to cluster timeseries into brain states; thus, we focused more on the temporal rather than spatial resolution of the data. Under no drug influence, Kay et al. (2012) reported similar patterns of DMN connectivity in subjects who were listening to music compared with those who were not. These results are similar to our observations of brain states’ dynamics regarding the DMN activity which does not differ between resting-state and music, except that we also found the lack of significant effect after the LSD intake.

It is important to note that previous studies revealed that psychedelic experience combined with musical stimuli has a profound influence on the subject’s mental condition, especially emotion processing (Carbonaro et al., 2018; Kaelen et al., 2015; Kaelen, Roseman, Lorenz, et al., 2016) and meaning-making (Preller et al., 2017). Here, we hypothesized that such a strong impact on the psychological state should also be accompanied by the changes in the dynamics of the time-varying brain activity. Specifically, we expected to observe different patterns of brain states’ dynamics for the resting-state *before* and *after* listening to music. We found that resting-state runs, separated by music listening, have similar brain states’ dynamics regarding the duration of states and their frequency, for both LSD and placebo sessions. Contrarily, we observed the difference between the two resting-states in terms of the states’ transitions. We found that on LSD, resting-state after music listening showed higher probability of transition from state associated with high DMN and FPN networks activity to state characterized by high FPN, DAN and VAN activity, in comparison to resting-state prior to musical stimuli. Additionally, the difference between both resting-state runs for this transition was significantly greater under LSD, what highlights the potential long-term effect of musical stimuli and suggests that LSD combined with music promotes transitions between states, while states’ fluctuations during placebo are more static and organized. Here, such pattern of transitions may reflect switching from common brain activity after psychedelics intake, associated with self-referential processing and mind-wandering, to activity crucial for cognitive control and integration of multiple stimuli; thus we may conclude that it could be crucial for processing the whole psychedelic experience. Previous studies showed that the joint effect of psychedelics and music affects brain activity and network architecture (Barrett, Preller, Herdener, et al., 2018; Kaelen, Roseman, Kahan, et al., 2016; Kaelen, Roseman, Lorenz, et al., 2016; Preller et al., 2017). In particular, Kaelen et al. (2016) reported increased functional connectivity between the parahippocampus and visual cortex, while Preller et al. (2017) showed greater BOLD activity in the SMA, putamen, insula, and the PCC. Together, our observations expand upon the previous studies by demonstrating that music combined with psychedelics may have a long-term effect on the resting-state that can be observed in the dynamics of the time-varying brain activity.

### Limitations

The main limitation of this study is the small sample size, which is a common problem for the studies involving psychedelics intake. The consequence of the small sample size is a low statistical power of the study, thus a high probability of the occurrence of false positives and false negatives. It is important to emphasize that these results should be considered as preliminary and further studies need to replicate the analysis and test our findings on larger samples. Additionally, for the brain states identification we used the K-Means algorithm, which is the most frequently used in terms of clustering fMRI timeseries. However, brain states can be also identified using other clustering methods, such as hierarchical clustering (Ward, 1963) or Hidden Markov-Model (Ghassempour et al., 2014; Li & Biswas, 1999). Performing the analysis of brain states’ dynamics using a different approach could test whether the effect of LSD and music is also present and consistent while applying other methodologies. Finally, we cannot exclude the possibility that the difference between the first and second resting-state scans do not depend on the drug effect reaching its peak intensity. Future studies should use more music-oriented study design to have better control over the effects of both psychedelics and stimuli.

## Conclusions

Here, we investigated the effect of psychedelics and music on the time-varying brain activity. We applied an unsupervised machine learning method KMeans to identify brain states and characterized their dynamics by calculating appropriate measures. Our study is the first to investigate an interaction between psychedelics and music on the brain states’ dynamics in the long-term. We show that musical stimuli, as a part of “setting”, can have an influence on the subject’s brain activity, and in consequence support the psychological outcomes of the psychedelics experience, however, further studies are needed to confirm these effects on the larger sample size.

## Acknowledgments

Project was funded by Nicolaus Copernicus University grant Grants4NCUStudents (2506). We also thank Ondrej Zika, PhD for his advice on conducting the statistical analysis.

