## Supplementary Information for "Effect of LSD and music on the time-varying brain dynamics"

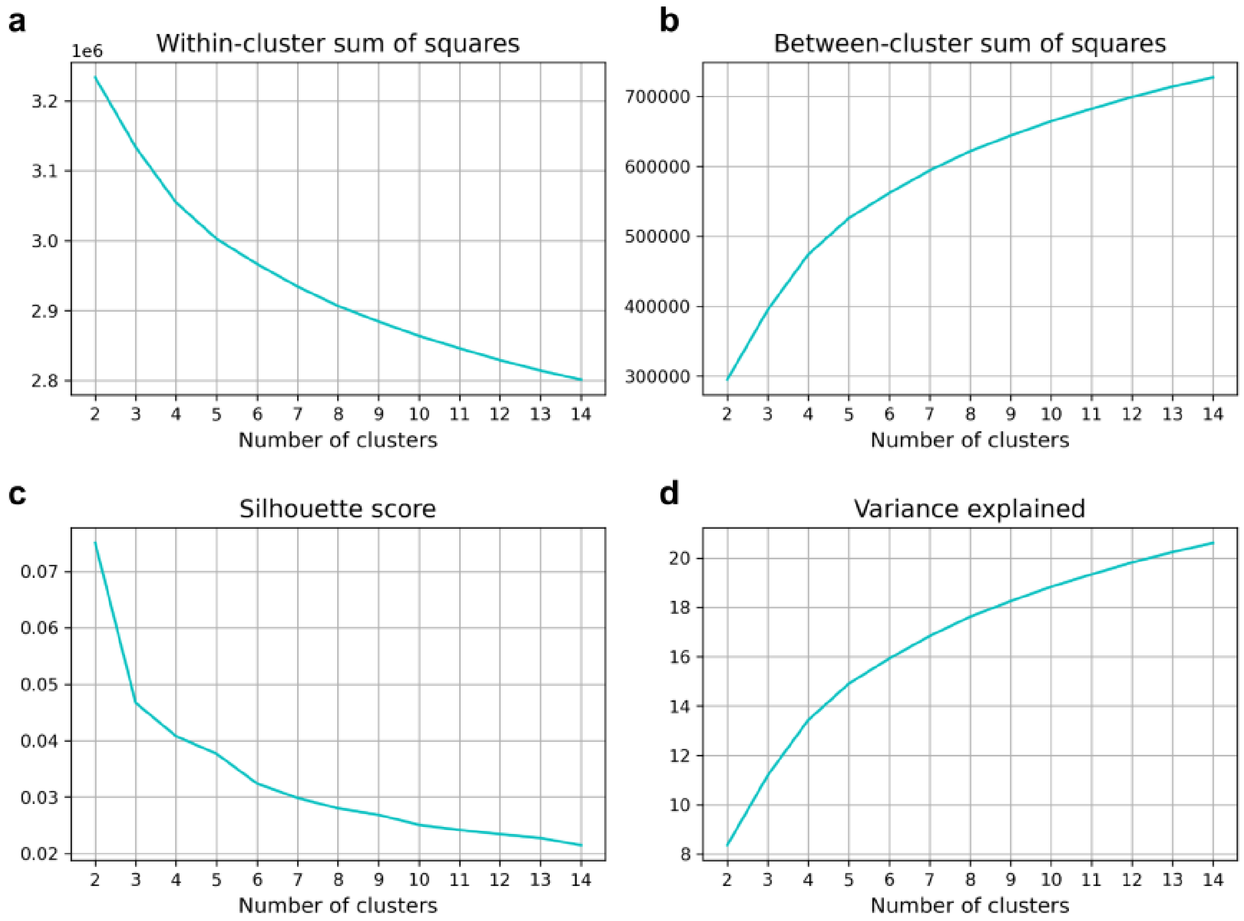


**Supplementary Fig. 1** K-Means measures calculated for *k* ranging from 2 to 14. **a** Within-cluster sum of squares, defined as the distances of each data point in all clusters to their respective centroids. **b** Between-cluster sum of squares, that is the squared average distance between the all centroids. **c** Silhouette score, which can be understood as a measure of how similar an object is to its own cluster compared to other clusters. **d** Variance explained, defined as the ratio of between-cluster sum of squares to the total sum of squares multiplied by 100.

**a**

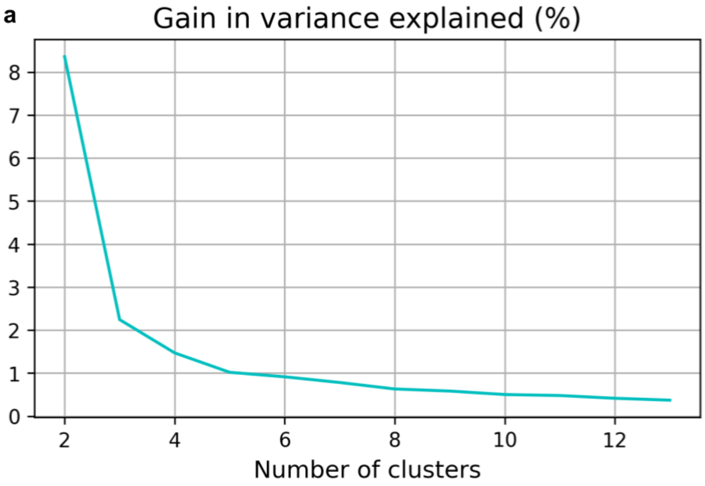


**b**

| k | gain in variance explained |
|---|---|
| 2 | 8.3564 |
| 3 | 2.24456 |
| 4 | 1.4711 |
| 5 | 1.0195 |
| 6 | 0.9148 |
| 7 | 0.7826 |
| 8 | 0.6320 |
| 9 | 0.5836 |
| 10 | 0.5029 |
| 11 | 0.4800 |
| 12 | 0.4157 |
| 13 | 0.3721 |

**Supplementary Fig. 2** Gain in variance explained. **a** Plot visualizing gain in variance explained for *k* ranging from 2 to 13. **b** Table with gain in variance explained values.

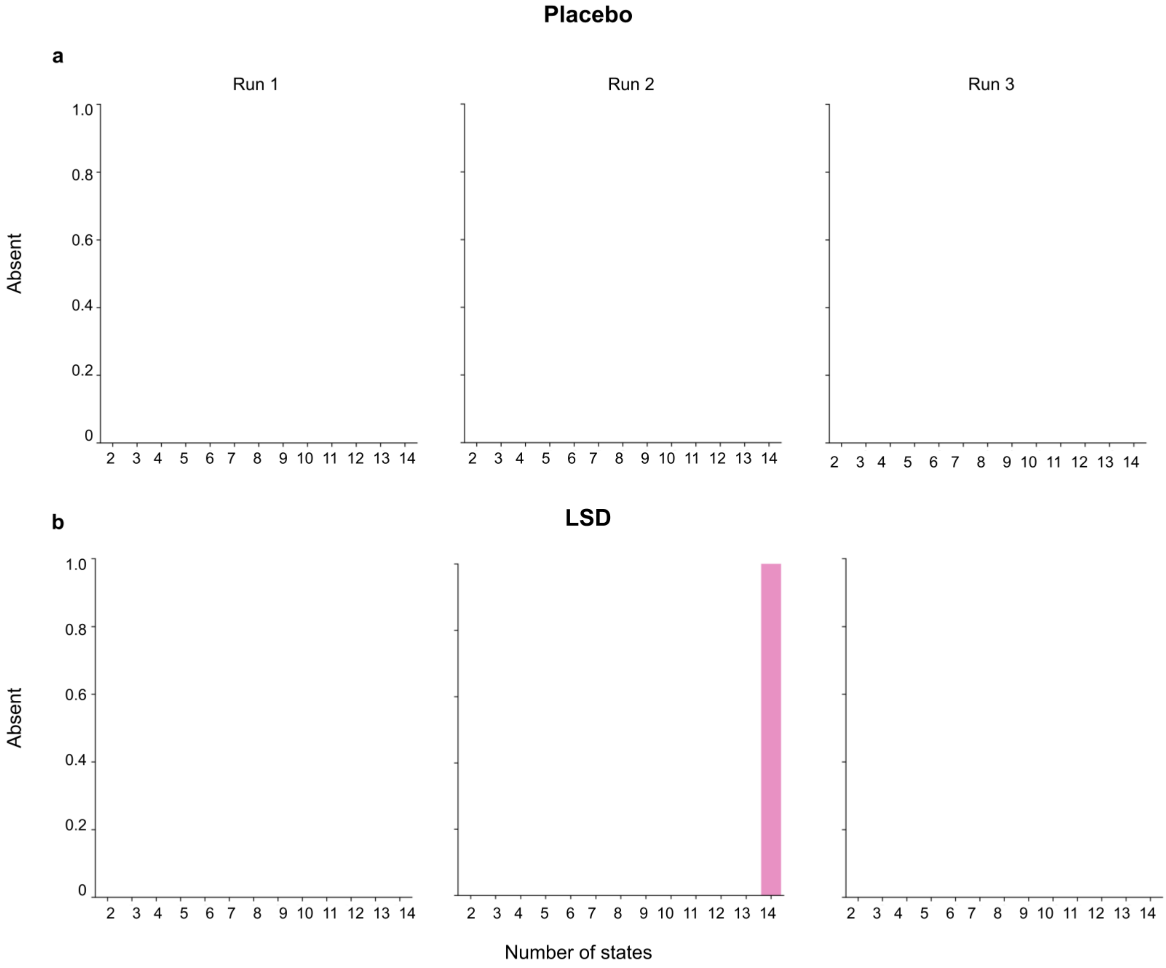


**Supplementary Fig. 3** Number of missing brain states for *k* ranging from 2 to 14. **a** The amount of absent states calculated for all runs for the placebo session. We found that for every *k* the correspondence of states between subjects was maintained. **b** The amount of absent states calculated for all runs for the LSD session. We observed that for $k > 13$ the correspondence of states across subjects was disturbed.

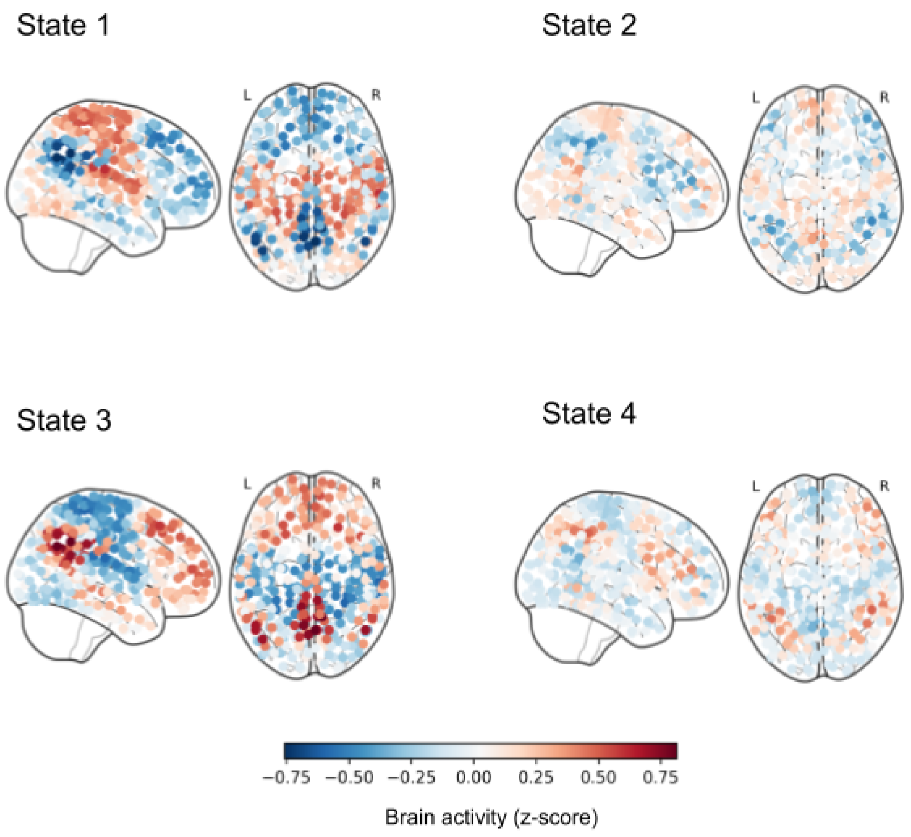


**Supplementary Fig. 4** Brain states' activation patterns for 4 states presented on glass brain plots.

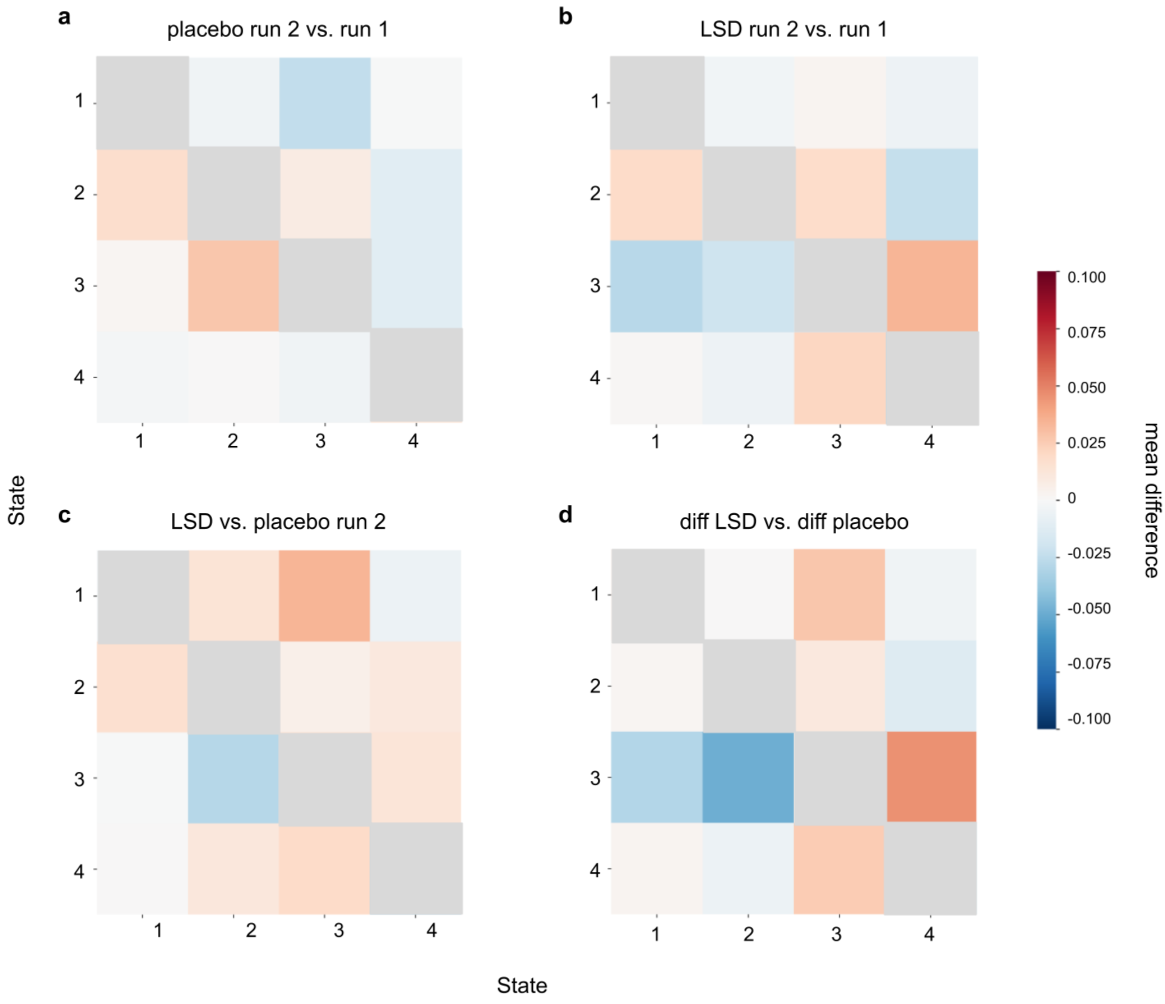


**Supplementary Fig. 5** Transition matrices for the comparison of resting-state and music experience. For all comparisons (a, b, c, d) the difference was not statistically significant. For all transition matrices, the direction of transition from one state to the other is row to column.

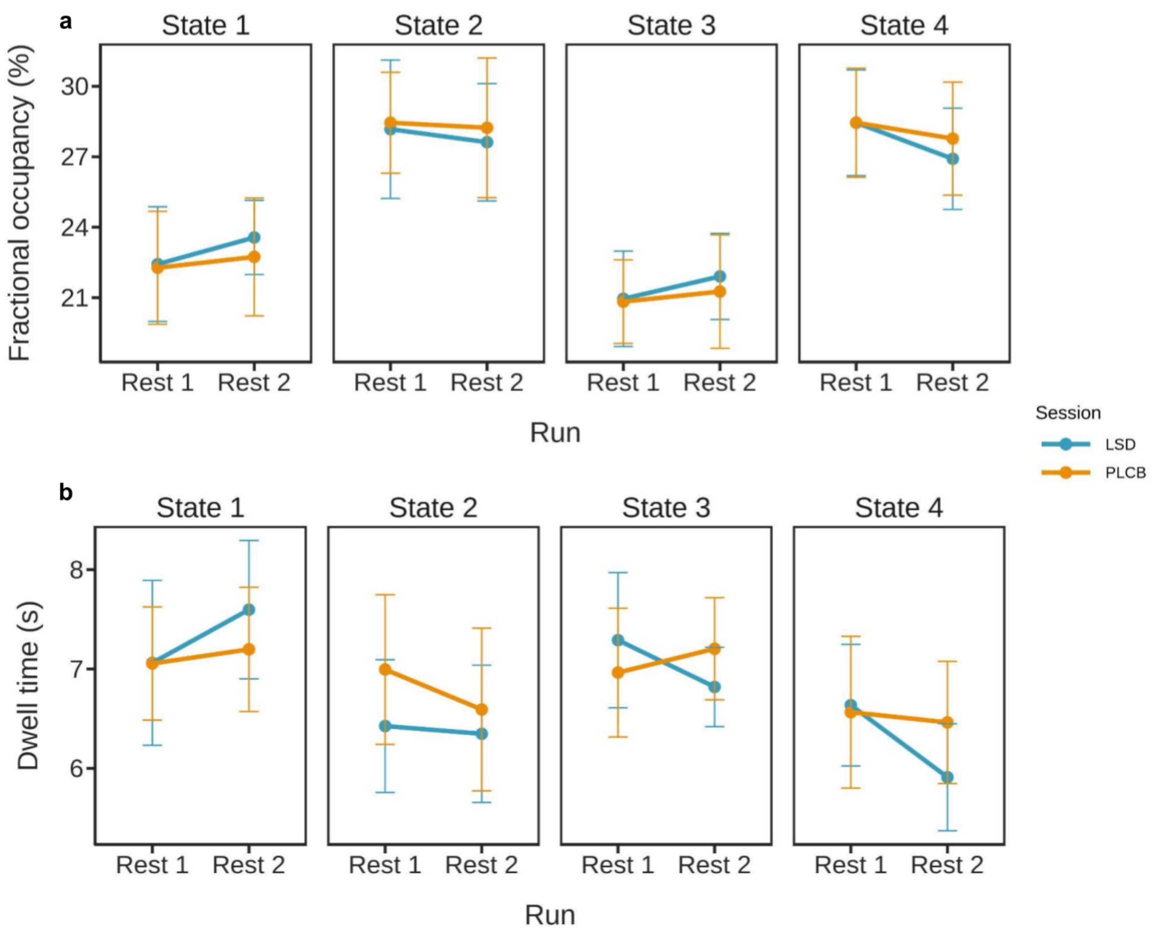


**Supplementary Fig. 6** Comparison of pre- and post-music listening resting-state. **a** Fractional occupancy for 1st (resting-state *before* music listening) and 3rd (resting-state *after* music listening) runs for both LSD and placebo conditions. For every state, we found no significant main and interaction effects. **b** Dwell time for 1st (resting-state *before* music listening) and 3rd (resting-state *after* music listening) runs for both LSD and placebo conditions. For every state, we found no significant main and interaction effects.

### Analysis for 5 states

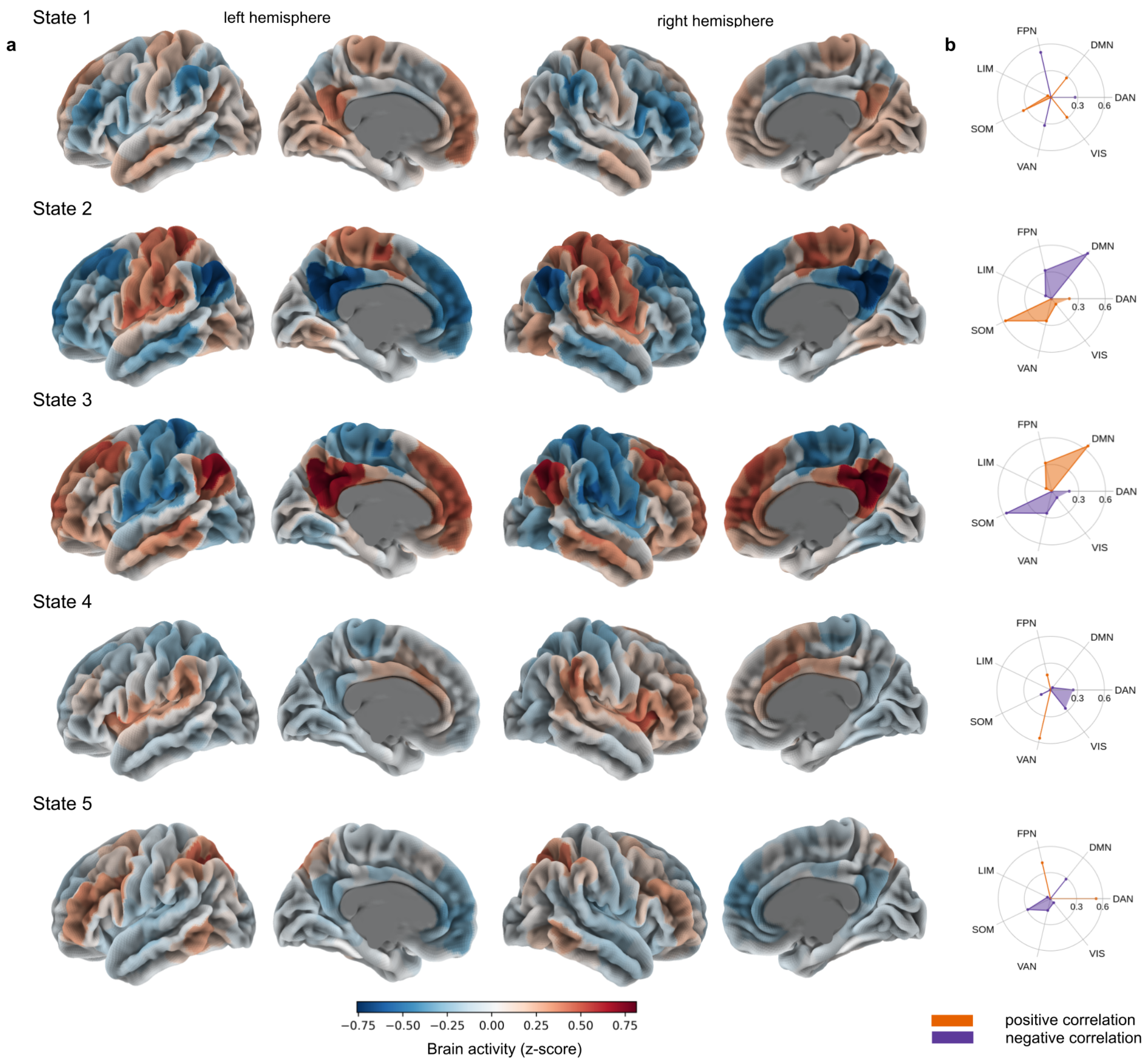


**Supplementary Fig. 7** Brain states' correlation with large-scale brain networks for 5 states. **a** Each state is defined as a characteristic pattern of brain activity, which fluctuates from high to low. **b** Correlation with 7 large-scale brain networks obtained from Schaefer 2018 parcellation (FPN - frontoparietal network, DMN - default mode network, DAN - dorsal attention network, VIS - visual network, VAN - ventral attention network, SOM - somatomotor network, LIM - limbic network).

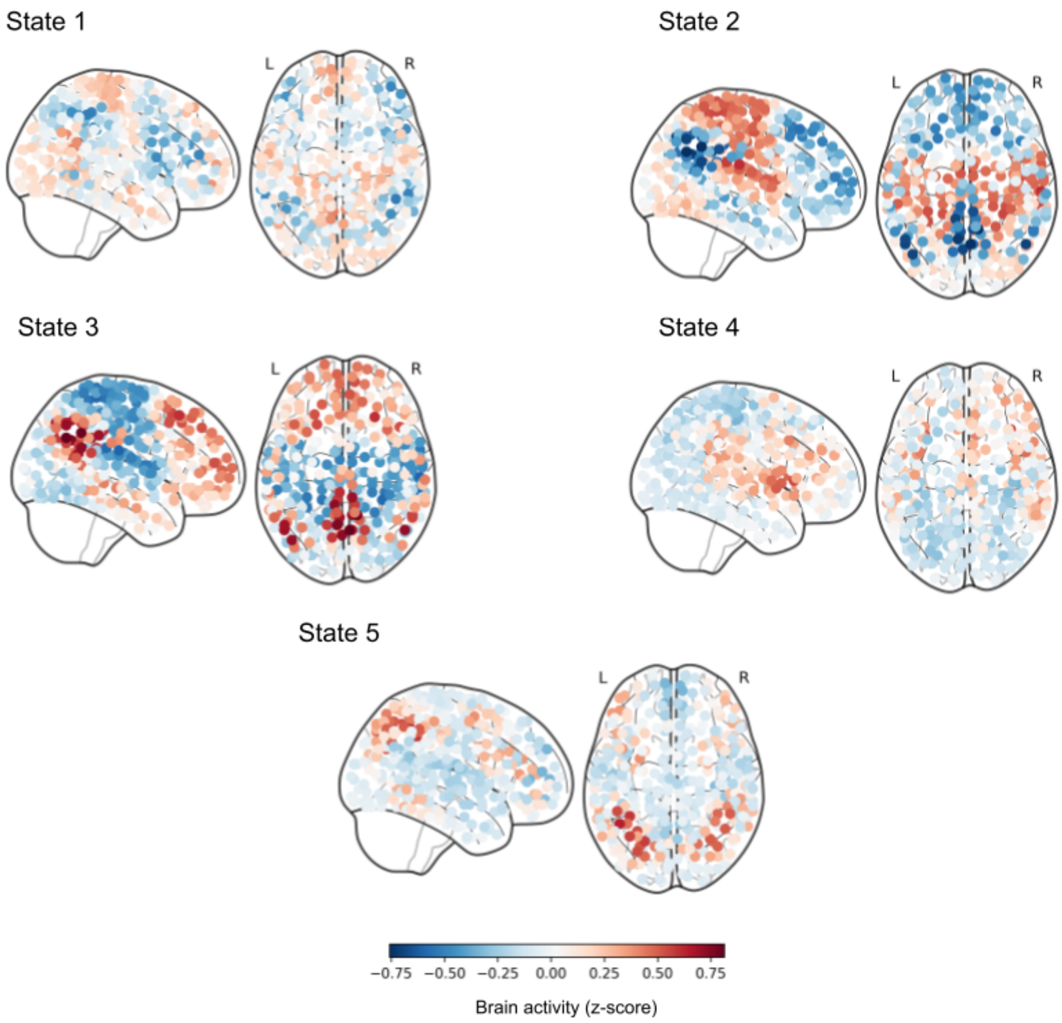


**Supplementary Fig. 8** Brain states' activation patterns for 5 states presented on glass brain plots.

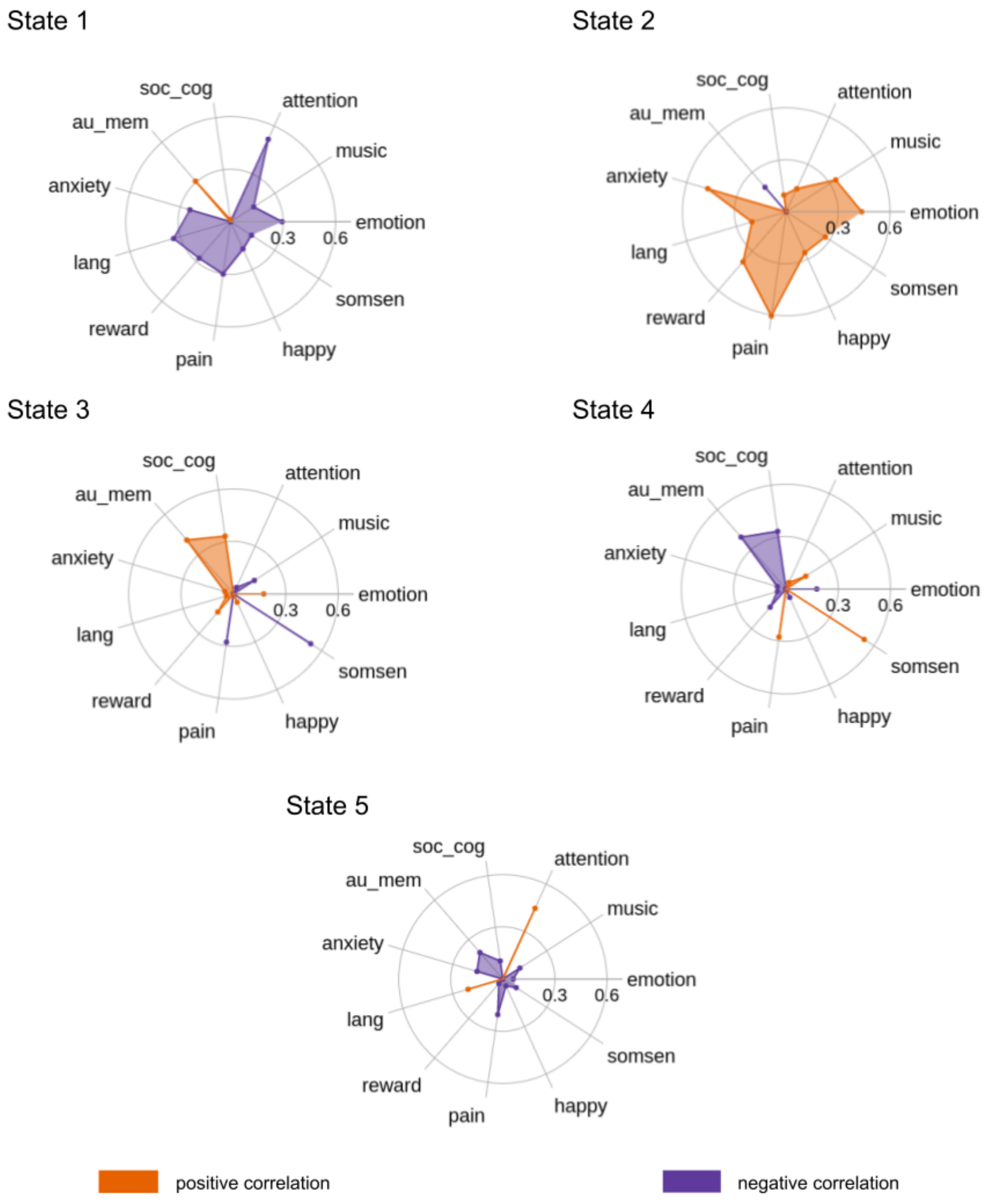


**Supplementary Fig. 9** Brain states' functional profiles for 5 states. Uniformity test maps obtained from the Neurosynth database can potentially describe each brain state in the terms of potentially associated mental states, which can occur during psychedelic experience (soc_cog - social cognition, emotion - emotional, somsen - somatosensory, lang - language, au_mem - autobiographical memory).

Effect of LSD on brain states during music experience

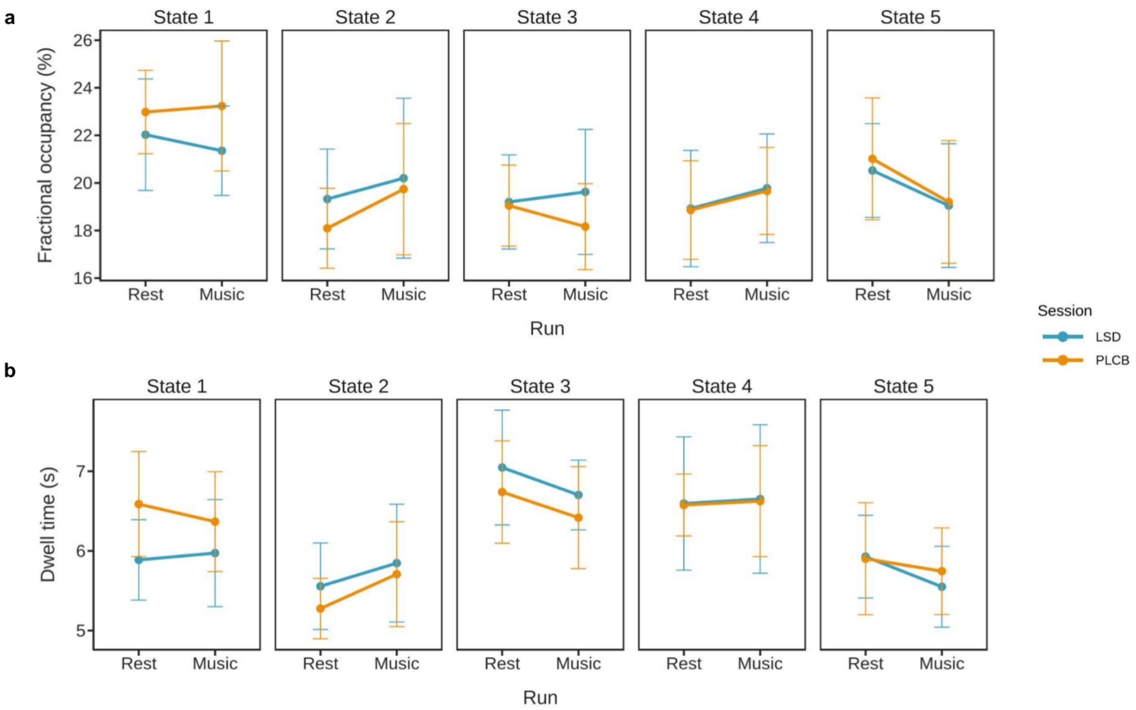


**Supplementary Fig. 10** Comparison of the resting-state and music experience for 5 states. **a** Fractional occupancy for 1st (resting-state) and 2nd (music listening) runs for both LSD and placebo conditions. For every state, we found no significant main and interaction effects. **b** Dwell time for 1st (resting-state) and 2nd (music listening) runs for both LSD and placebo conditions. For every state, we found no significant main and interaction effects.

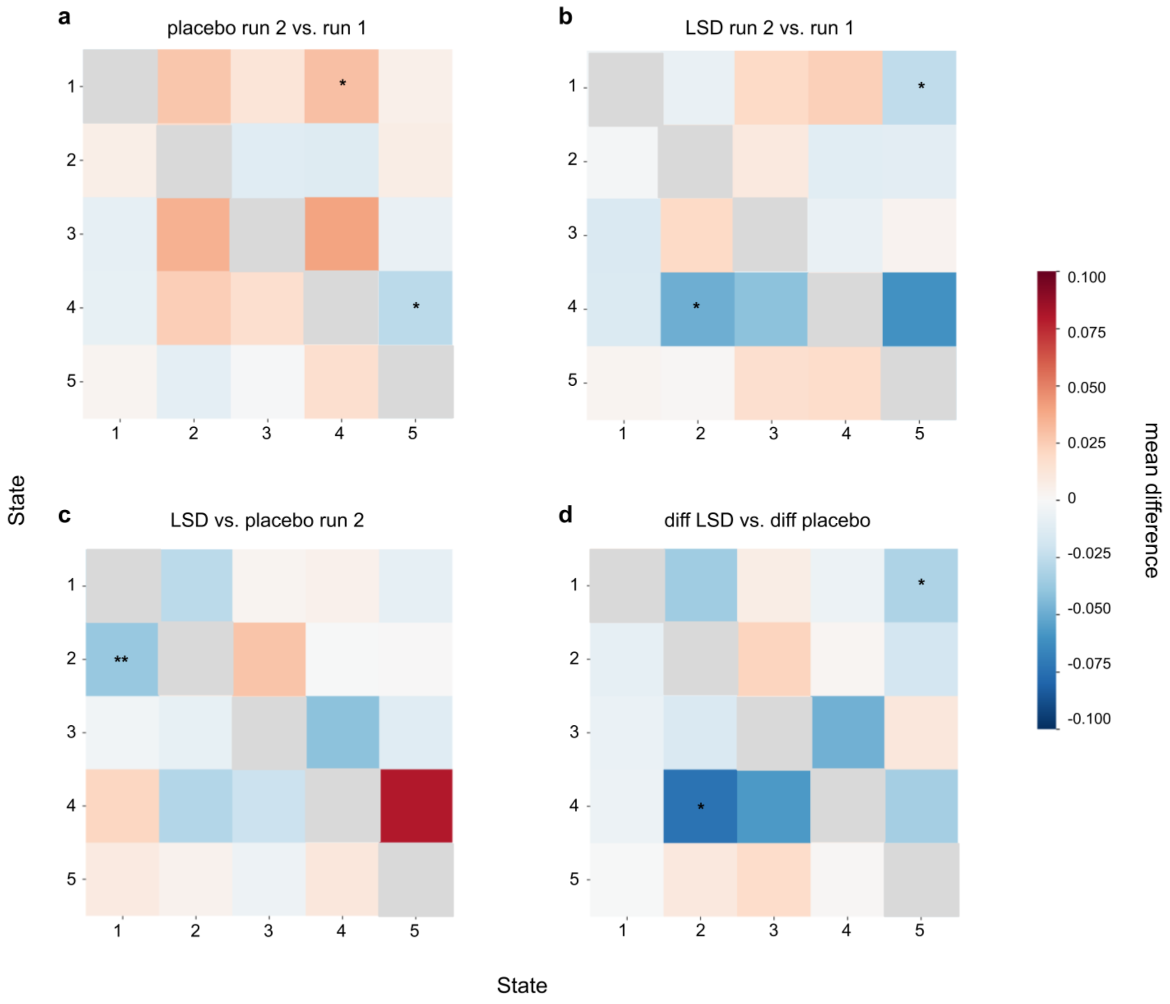


**Supplementary Fig. 11** Comparison of the resting-state and music experience transition patterns for 5 states. **a** Music listening on placebo was associated with with higher probability of transitioning from state 1 to state 4 (permutation test, mean difference = 0.03, $p$ = 0.015) and lower probability of transitioning from state 4 to state 5 (permutation test, mean difference = -0.027, $p$ = 0.034) than during resting state. **b** Transition matrix for the LSD session, 2rd run vs. 1st run; we found that music listening on LSD was associated with lower probability of transitioning from state 4 to 2 (permutation test, mean difference = -0.049, $p$ = 0.046) and from state 1 to 5 (permutation test, mean difference = -0.025, $p$ = 0.01) in comparison to resting state. **c** During LSD session in comparison with placebo, music listening was associated with lower probability of transitioning from state 2 to state 1 (permutation test, mean difference = -0.038, $p$ = 0.0024). **d** Transition matrix for the differences (2rd run - 1st run) between both sessions (LSD and placebo). We found that for transition from state 4 to 2 (permutation test, mean difference = -0.074, $p$ = 0.035) and from state 1 to 5 (permutation test, mean difference = -0.031, $p$ = 0.045) the difference between both runs was significantly lower after the LSD intake. For all transition matrices, the direction of transition from one state to the other is row to column.

Effect of LSD and music listening on brain states during resting-state

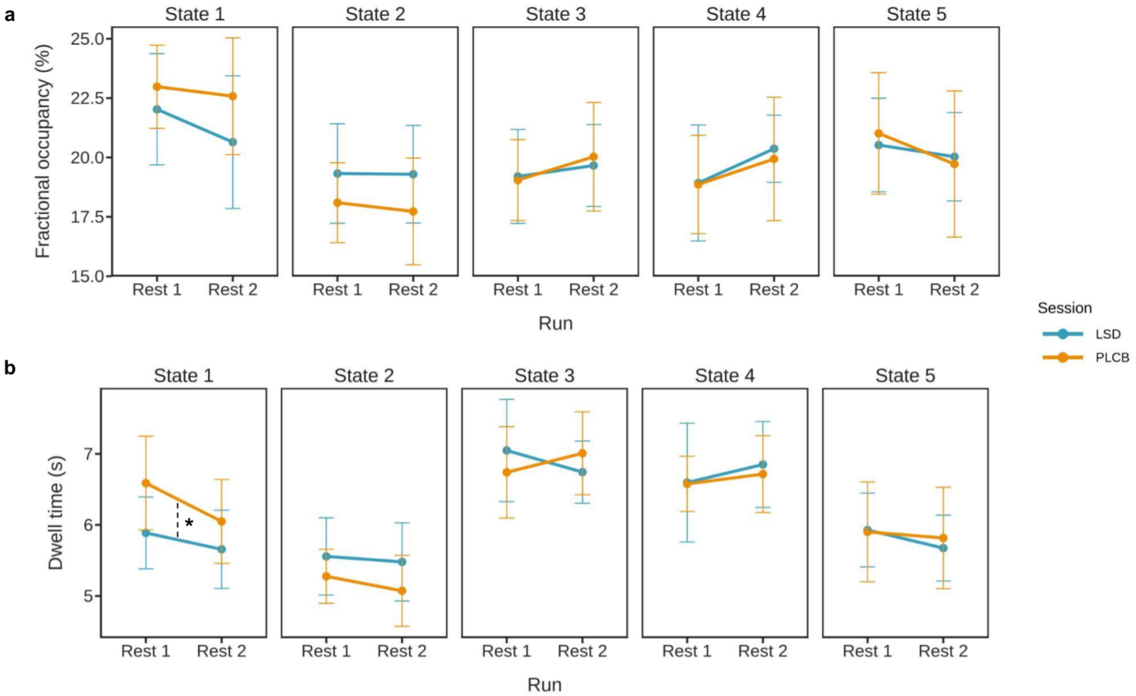


**Supplementary Fig. 12** Comparison of the pre- and post-music listening resting-state for 5 states. **a** Fractional occupancy for 1st (resting-state *before* music listening) and 3rd (resting-state *after* music listening) runs for both LSD and placebo conditions. For every state, we found no significant main and interaction effects. **b** Dwell time for 1st (resting-state *before* music listening) and 3rd (resting-state *after* music listening) runs for both LSD and placebo conditions. We found a significant session effect for state 1; during placebo session the dwell time of this state was significantly higher in comparison with LSD ($F_{1,42} = 4.8748$, $p = 0.033$).

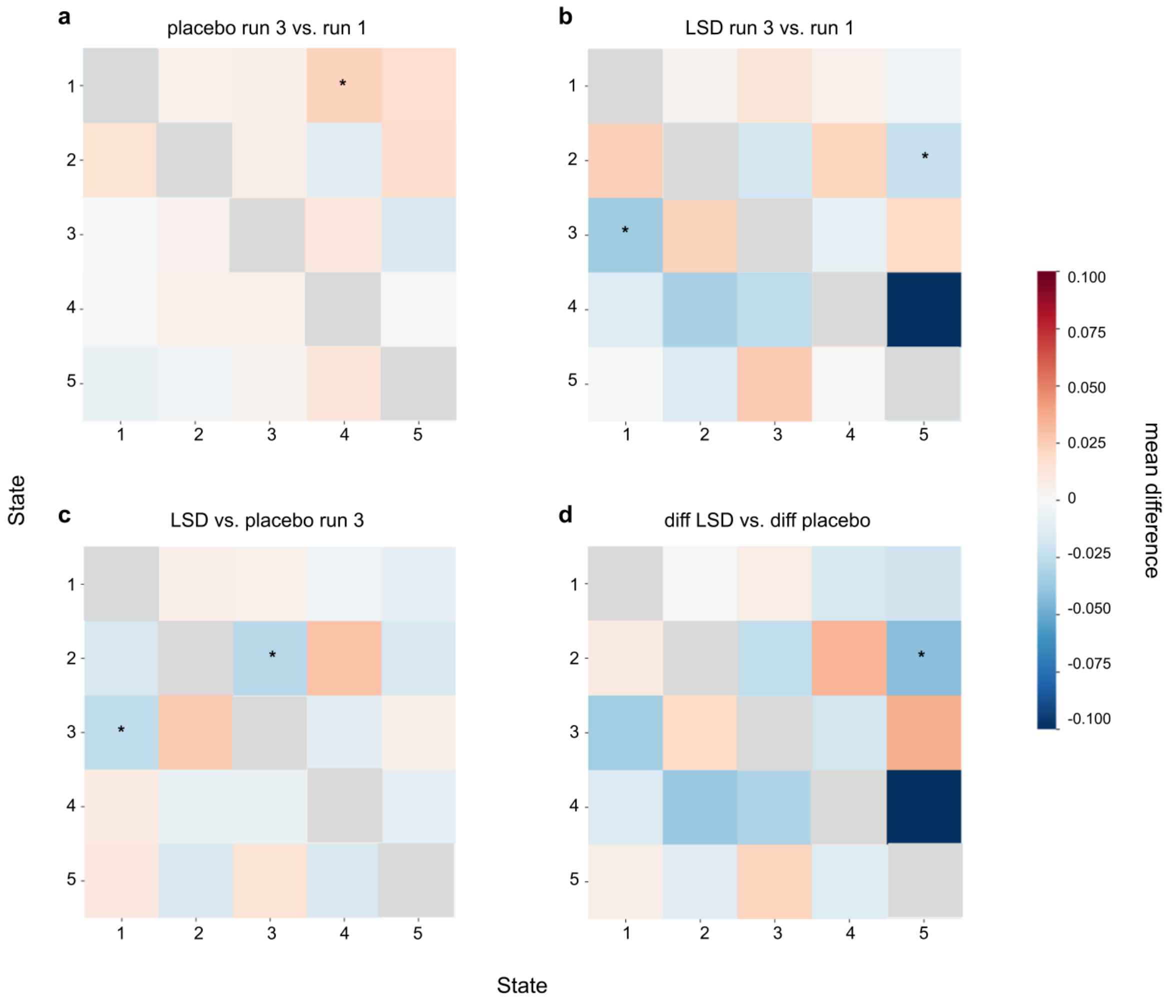


**Supplementary Fig. 13** Comparison of the pre- and post-music listening resting-state for 5 states. **a** During placebo, the resting-state after listening to music was associated with higher probability of transitioning from state 1 to 4 (permutation test, mean difference = 0.023, $p$ = 0.028). **b** Transition matrix for the LSD session, 3rd run vs. 1st run; under LSD influence, run 3 was associated with lower probability of transitioning from state 3 to 1 (permutation test, mean difference = -0.035, $p$ = 0.038) as well as from state 2 to 5 (permutation test, mean difference = -0.024, $p$ = 0.048). **c** During LSD session in comparison with placebo, resting-state after listening to music was associated with lower probability of transitioning from state 3 to state 1 (permutation test, mean difference = -0.025, $p$ = 0.049) and from state 2 to 3 (permutation test, mean difference = -0.028, $p$ = 0.037). **d** Transition matrix for the differences (3rd run - 1st run) between both sessions (LSD and placebo). The results show that under the LSD influence there is a less change in transition probability for transitions from state 2 to 5 (permutation test, mean difference = -0.042, $p$ = 0.034) in comparison with placebo. For all transition matrices, the direction of transition from one state to the other is row to column.

#### Analysis for 6 states

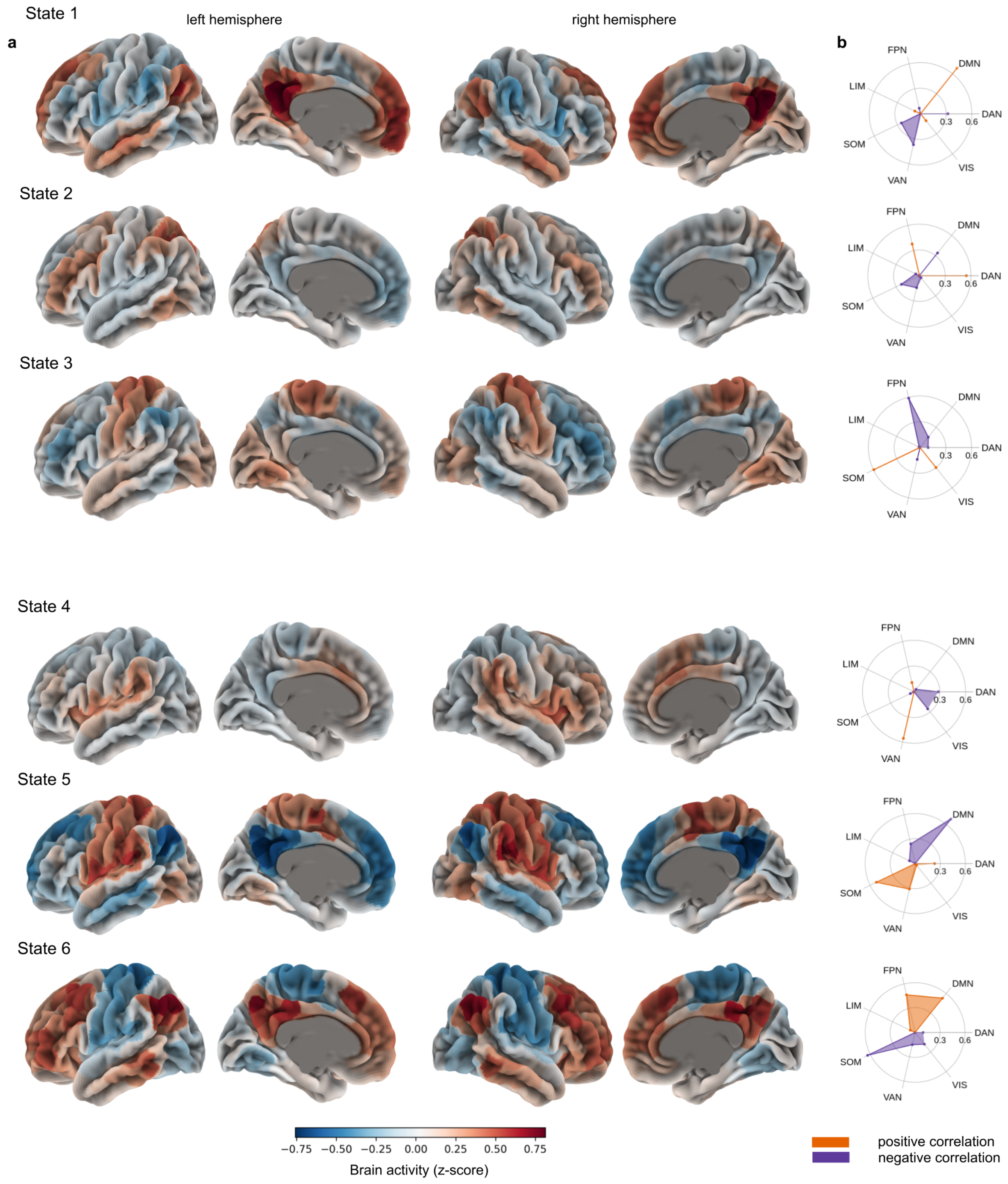


**Supplementary Fig. 14** Brain states' correlation with large-scale brain networks for 6 states. **a** Each state is defined as a characteristic pattern of brain activity, which fluctuates from high to low. **b** Correlation with 7 large-scale brain networks obtained from Schaefer 2018 parcellation (FPN - frontoparietal network, DMN - default mode network, DAN - dorsal attention network, VIS - visual network, VAN - ventral attention network, SOM - somatomotor network, LIM - limbic network).

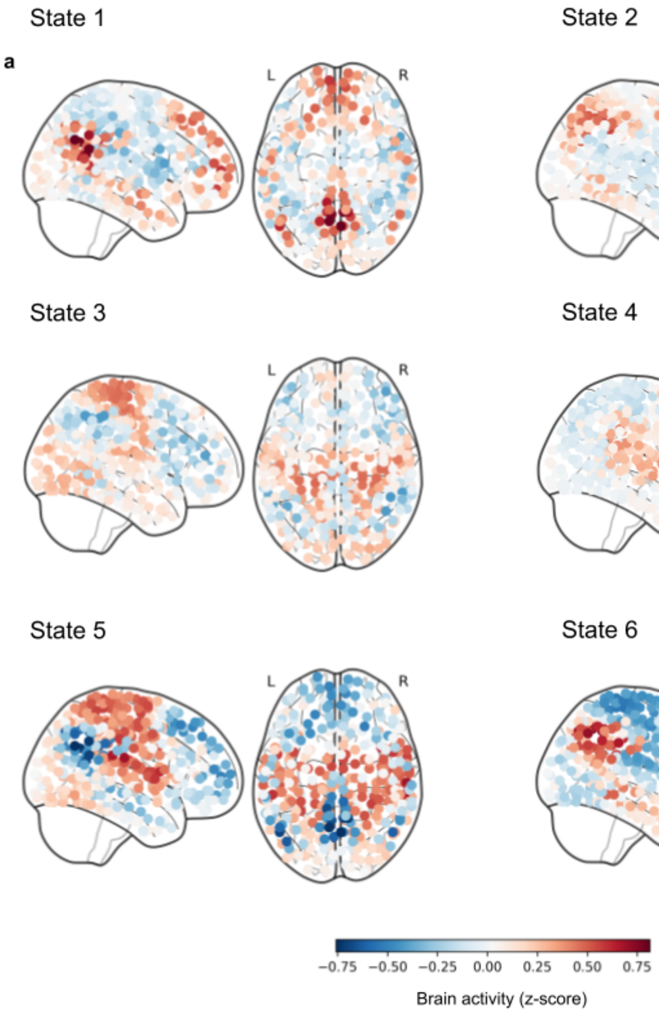


**Supplementary Fig. 15** Brain states' activation patterns for 6 states presented on glass brain plots.

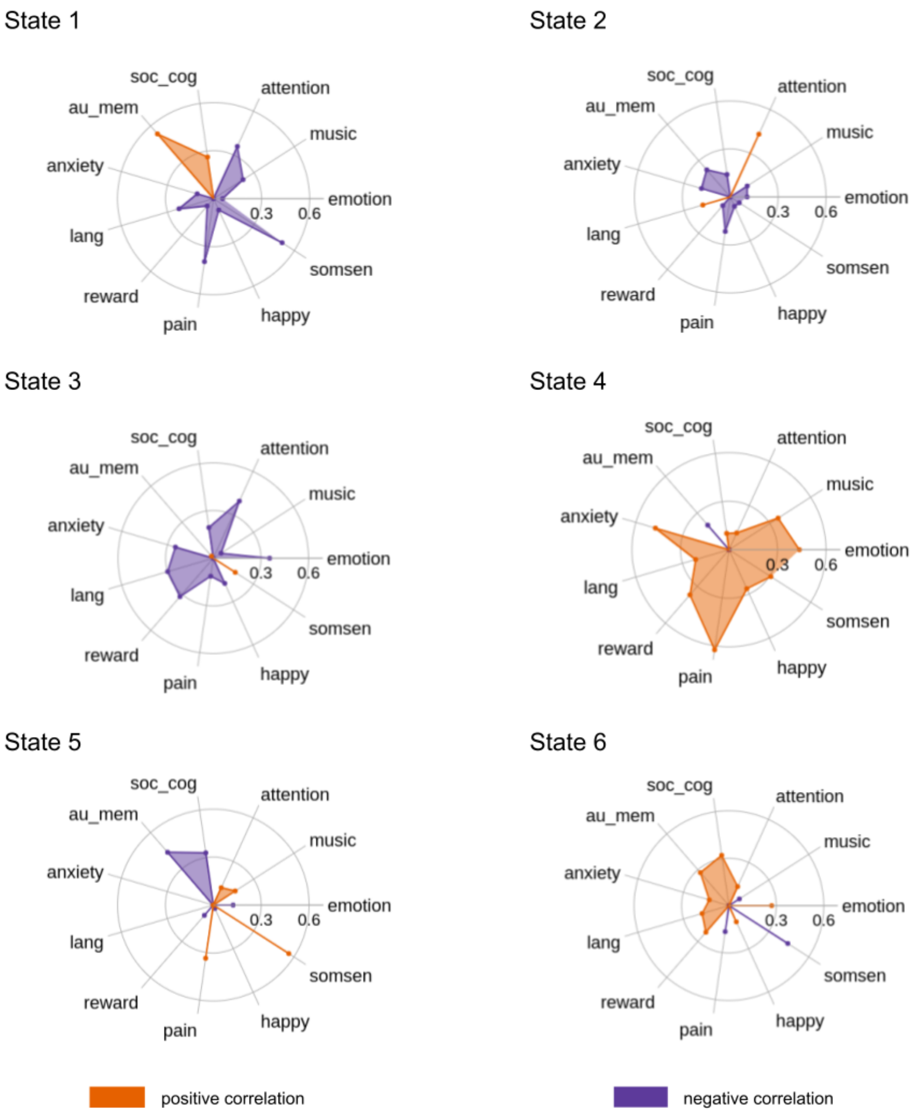


**Supplementary Fig. 16** Brain states' functional profiles for 6 states. Uniformity test maps obtained from the Neurosynth database can potentially describe each brain state in the terms of potentially associated mental states, which can occur during psychedelic experience (soc_cog - social cognition, emotion - emotional, somsen - somatosensory, lang - language, au_mem - autobiographical memory).

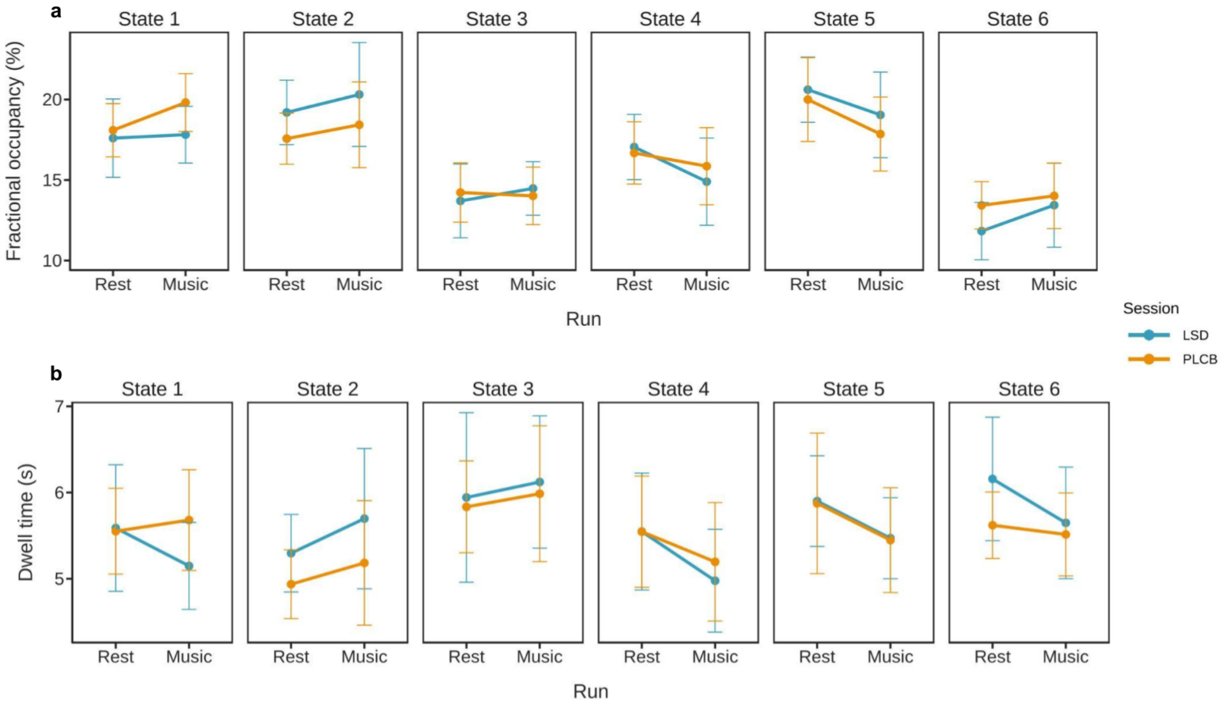


**Supplementary Fig. 17** Comparison of the resting-state and music experience for 6 states. **a** Fractional occupancy for 1st (resting-state) and 2nd (music listening) runs for both LSD and placebo conditions. For every state, we found no significant main and interaction effects. **b** Dwell time for 1st (resting-state) and 2nd (music listening) runs for both LSD and placebo conditions. For every state, we found no significant main and interaction effects.

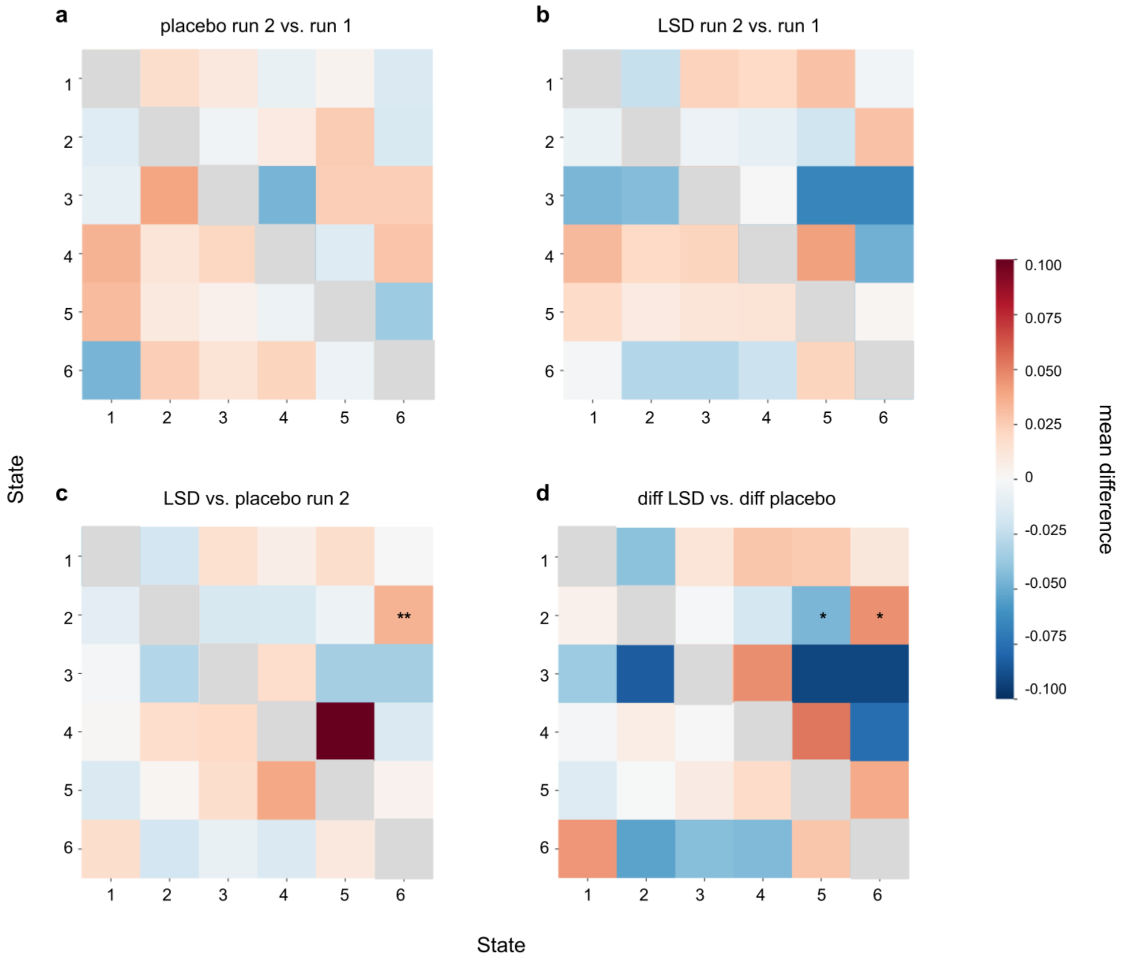


**Supplementary Fig. 18** Comparison of the resting-state and music experience transition patterns for 6 states. **a** Transition matrix for the placebo session, 2rd run vs. 1st run; we found no significant differences. **b** Transition matrix for the LSD session, 2rd run vs. 1st run; we found no significant differences. **c** During LSD session in comparison with placebo, music listening was associated with higher probability of transitioning from state 2 to state 6 (permutation test, mean difference = 0.035, $p$ = 0.0022) **d** Transition matrix for the differences (2rd run - 1st run) between both sessions (LSD and placebo). We found that for transition from state 2 to 5 the difference between both runs was significantly lower after the LSD intake (permutation test, mean difference = -0.046, $p$ = 0.015). In contrast, for transition from state 2 to 6 the difference was greater for the LSD session (permutation test, mean difference = 0.45, $p$ = 0.021). For all transition matrices, the direction of transition from one state to the other is row to column.

Effect of LSD and music listening on brain states during resting-state

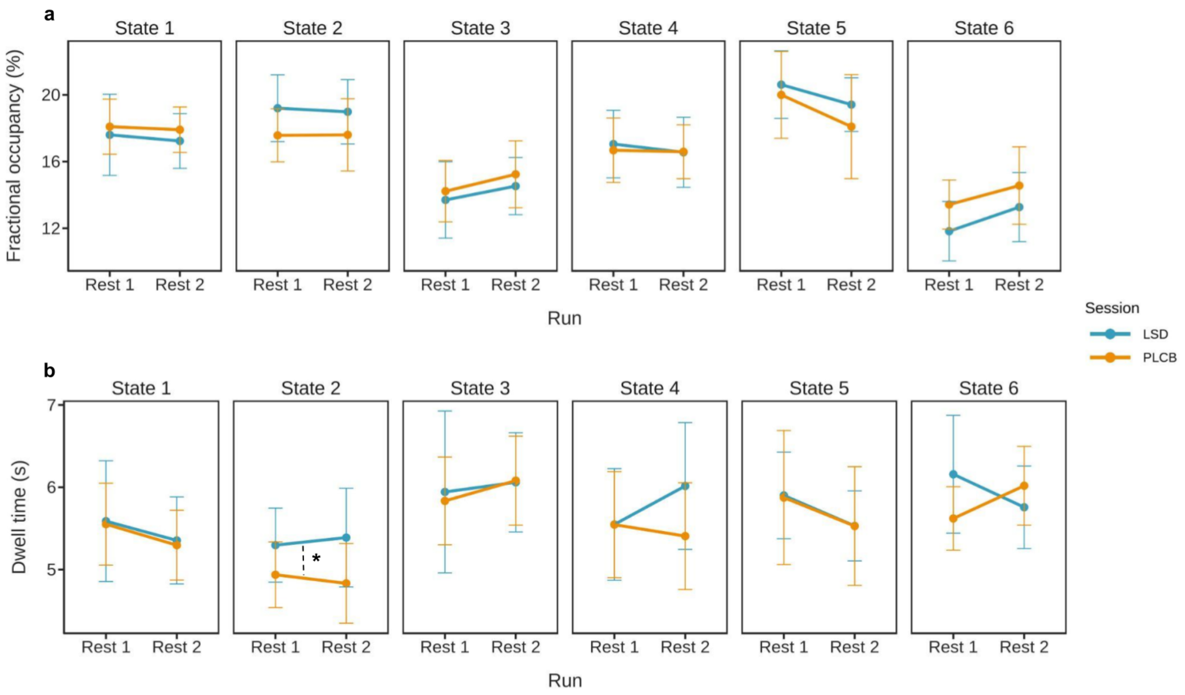


**Supplementary Fig. 19** Comparison of the pre- and post-music listening resting-state for 6 states. **a** Fractional occupancy for 1st (resting-state *before* music listening) and 3rd (resting-state *after* music listening) runs for both LSD and placebo conditions. For every state, we found no significant main and interaction effects. **b** Dwell time for 1st (resting-state *before* music listening) and 3rd (resting-state *after* music listening) runs for both LSD and placebo conditions. We found a significant session effect for state 2; during LSD session the dwell time of this state was significantly higher in comparison with placebo ($F_{1,42} = 4.3518$, $p = 0.043$).

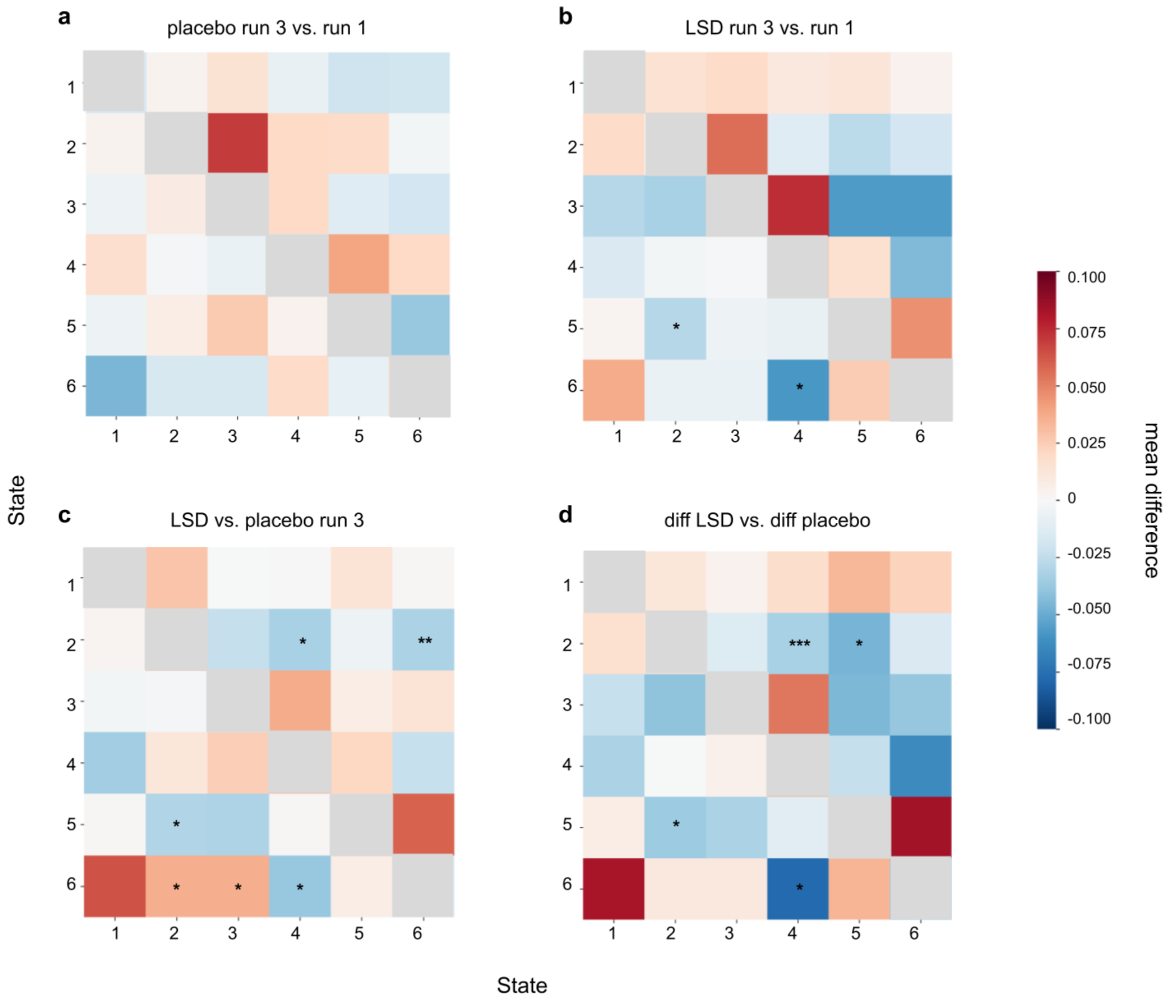


**Supplementary Fig. 20** Comparison of pre- and post-music listening resting-state for 6 states. **a** Transition matrix for the placebo session, 3rd run vs. 1st run; we found no significant differences. **b** Transition matrix for the LSD session, 3rd run vs. 1st run; under LSD influence, run 3 was associated with lower probability of transitioning from state 5 to 2 (permutation test, mean difference = -0.029, $p$ = 0.03) and 6 to 4 (permutation test, mean difference = -0.059, $p$ = 0.02). **c** During LSD session in comparison with placebo, resting-state after listening to music was associated with higher probability of transitioning from state 6 to state 2 (permutation test, mean difference = 0.037, $p$ = 0.028) and 3 (permutation test, mean difference = 0.037, $p$ = 0.028). Contrarily, in this condition there was lower probability of transitioning from state 5 to 2 (permutation test, mean difference = -0.03, $p$ = 0.012), from state 2 to 4 (permutation test, mean difference = -0.032, $p$ = 0.023) and 6 to 4 (permutation test, mean difference = -0.038, $p$ = 0.041) as well as from state 2 to 6 (permutation test, mean difference = -0.032, $p$ = 0.0084. **d** Transition matrix for the differences (3rd run - 1st run) between both sessions (LSD and placebo). The results show that under the LSD influence there is a less change in transition probability for transitions from state 5 to 2 (permutation test, mean difference = -0.036, $p$ = 0.03), from state 2 to 4 (permutation test, mean difference = -0.032, $p$ = 0.0009), from state 6 to 4 (permutation test, mean difference = -0.078, $p$ = 0.011) and from state 2 to 5 (permutation test, mean difference = -0.046, $p$ = 0.041). For all transition matrices, the direction of transition from one state to the other is row to column.
